# Dense connectomic reconstruction in layer 4 of the somatosensory cortex

**DOI:** 10.1101/460618

**Authors:** Alessandro Motta, Manuel Berning, Kevin M. Boergens, Benedikt Staffler, Marcel Beining, Sahil Loomba, Christian Schramm, Philipp Hennig, Heiko Wissler, Moritz Helmstaedter

**Author notes:** equally contributing first authors (Alessandro Motta, Manuel Berning, Kevin M. Boergens, Benedikt Staffler).

## Abstract

The dense circuit structure of the mammalian cerebral cortex is still unknown. With developments in 3-dimensional (3D) electron microscopy, the imaging of sizeable volumes of neuropil has become possible, but dense reconstruction of connectomes from such image data is the limiting step. Here, we report the dense reconstruction of a volume of about 500,000 μm^3^ from layer 4 of mouse barrel cortex, about 300 times larger than previous dense reconstructions from the mammalian cerebral cortex. Using a novel reconstruction technique, FocusEM, we were able to reconstruct a total of 0.9 meters of dendrites and about 1.8 meters of axons investing only about 4,000 human work hours, about 10-25 times more efficient than previous dense circuit reconstructions. We find that connectomic data alone allows the definition of inhibitory axon types that show established principles of synaptic specificity for subcellular postsynaptic compartments. We find that also a fraction of excitatory axons exhibit such subcellular target specificity. Only about 35% of inhibitory and 55% of excitatory synaptic subcellular innervation can be predicted from the geometrical availability of membrane surface, revoking coarser models of random wiring for synaptic connections in cortical layer 4. We furthermore find evidence for enhanced variability of synaptic input composition between neurons at the level of primary dendrites in cortical layer 4. Finally, we obtain evidence for Hebbian synaptic weight adaptation in at least 24% of connections; at least 35% of connections show no sign of such previous plasticity. Together, these results establish an approach to connectomic phenotyping of local dense neuronal circuitry in the mammalian cortex.

## INTRODUCTION

The cerebral cortex of mammals houses an enormously complex intercellular interaction network implemented via neuronal processes that are long and thin, branching, and extremely densely packed. Early estimates reported an expected 4 kilometers of axons and 400 meters of dendrites compressed into a cubic millimeter of cortical tissue (Braitenberg and Schüz, 1998). This high packing density of cellular processes has made the locally dense mapping of neuronal networks in the cerebral cortex challenging.

So far, reconstructions of cortical tissue have been either sparse (e.g., (da Costa and Martin, 2009; Han et al., 2018; Lee et al., 2016; Lubke et al., 2003; Oberlaender et al., 2011; Schmidt et al., 2017) or restricted to small volumes of about 1500 μm^3^(Kasthuri et al., 2015). Consequently the detailed network architecture of the cerebral cortex is unknown; in particular the question to what degree local neuronal circuits are explainable by geometric rules alone, or whether neurons exhibit innervation specificities beyond such geometric preferences is still debated (Kasthuri et al., 2015; Ko et al., 2011; Lee et al., 2016; Markram et al., 2015; Mishchenko et al., 2010). Here we report a dense reconstruction of local cortical tissue sized about 500,000 μm^3^, i.e. about 300 times larger than previous dense cortical reconstructions (Kasthuri et al., 2015).

We developed a dense reconstruction method, FocusEM, to obtain the reconstruction of about 2.7 meters of neurite (1.8 meters of axons and about 0.9 meters of dendrites) with an investment of about 4,000 human work hours. When compared to previous dense connectomic reconstructions, this constitutes an advance of about 10-fold (compared to dense reconstructions in the fly larva, (Eichler et al., 2017)), about 20-fold (cf. mouse retina (Helmstaedter et al., 2013)) and about 25-fold (cf. mouse cortex (Kasthuri et al., 2015)).

When analyzing the connectivity between 6,979 axons and 3,719 postsynaptic neurites in this tissue, we find that at least about 58% of inhibitory and about 24% of excitatory axons show specificity for synaptic targets such as cell bodies, apical dendrites and axon initial segments. We determine that only about 35-55% of this synaptic specificity can be deduced from the geometrical arrangement of axons and dendrites alone at scales typically employed for statistical connectivity prediction (Lubke et al., 2003; Meyer et al., 2010; Oberlaender et al., 2012a), establishing an upper bound on the geometrical explainability of synaptic innervation in cortical tissue. Furthermore, we find that the thalamocortical synaptic input distributions of dendrites are configured to yield enhanced variability. Finally, a fraction of excitatory axons show synaptic size similarity that is consistent with Hebbian plasticity, which at the same time can be ruled out for about 35% of the circuit. With this we uncover rules of wiring and synaptic specialization in the cerebral cortex and provide a methodology for connectomic screening of cortical tissue.

## RESULTS

We acquired a 3-dimensional EM dataset from a 28 day old mouse from layer 4 of primary somatosensory cortex (Fig. 1a-d) using serial block-face electron microscopy (SBEM, (Denk and Horstmann, 2004)). The dataset had a size of 61.8 × 94.8 × 92.6 μm^3^ and a voxel size of 11.24 × 11.24 × 28 nm^3^. We 3D-aligned the acquired images for manual annotation (webKnossos (Boergens et al., 2017)) and automated analysis. We first detected blood vessels and cell bodies using automated heuristics (Fig. 1e), followed by reconstruction of the remaining image volume using machine-learning-based image segmentation (SegEM, (Berning et al., 2015)). The result of this processing were 15 million volume segments corresponding to pieces of axons, dendrites and somata (volume: 0.0295 ± 0.3846 μm^3^; mean ± std.). We then constructed the neighborhood graph between all these volume segments and computed the properties of interfaces between directly adjacent volume segments. Based on these features (see Methods), we trained a connectivity classifier to determine whether two segments should be connected (along an axon or a dendrite or a glial cell) or whether they should be disconnected (Fig. 1f). Using the SynEM classifier (Staffler et al., 2017), we determined whether an interface between two disconnected processes corresponded to a chemical synapse, and if so, which was the pre-and which the postsynaptic neurite segment. We furthermore trained a set of classifiers to compute for each volume segment the probability to be part of an axon, a dendrite, a spine head or a glia cell (Fig. 1f-g, precision and recall were 91.8%, 92.9% for axons, 95.3%, 90.7% for dendrites, 97.2%, 85.9% for astrocytes, and 92.6%, 94.4% for spine heads, respectively).

**Fig.1.**
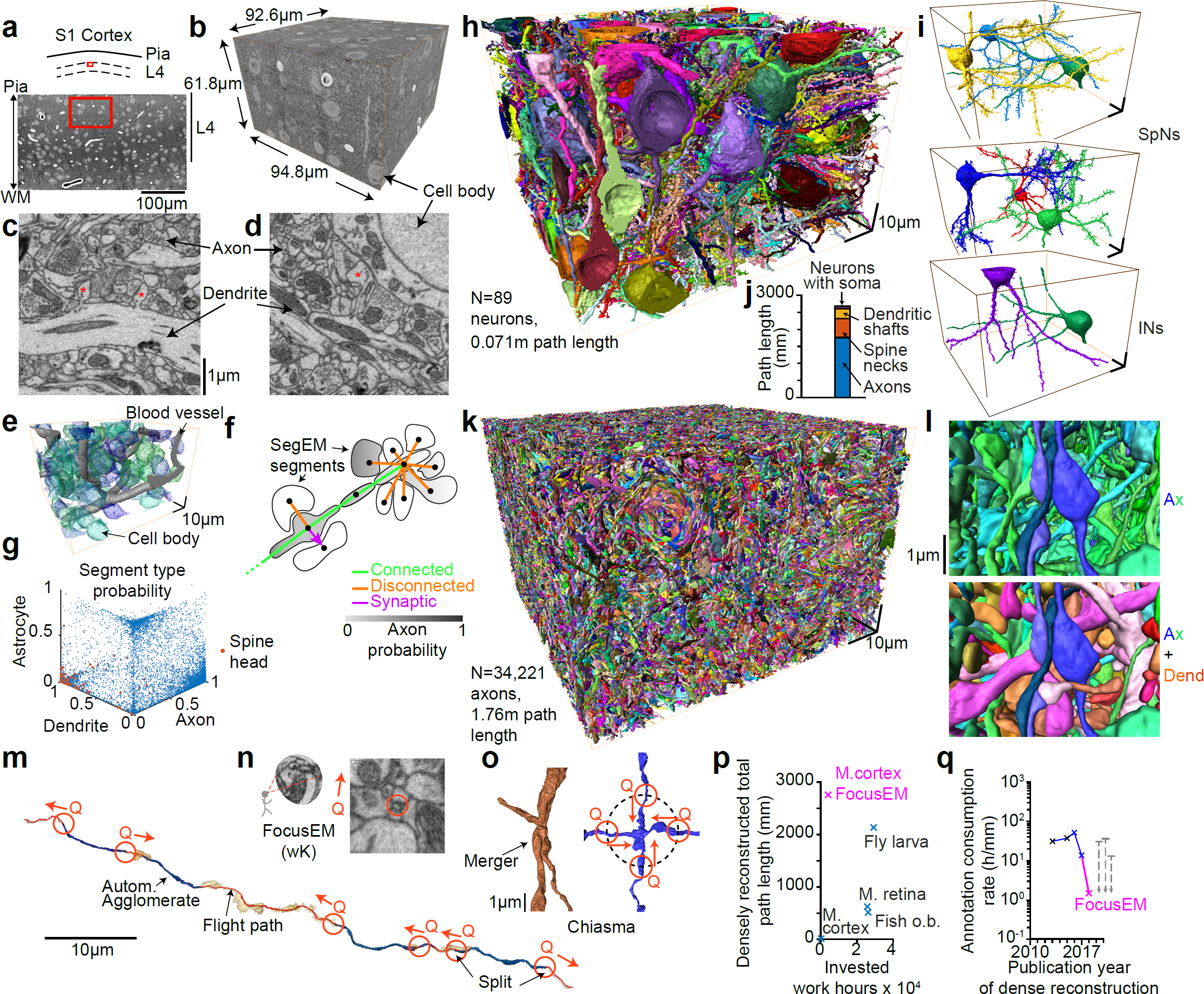
Dense connectomic reconstruction of cortical neuropil from layer 4 of mouse primary somatosensory cortex. (a-d) Location (a, red) of 3D EM dataset (b), WM: white matter; high-resolution example images (c,d). Asterisk, examples of dendritic spines. (e) Low-resolution automated reconstruction of cell bodies (blue-green) and blood vessels (gray). (f) Sketch of FocusEM, a set of methods for semi-automated reconstruction of dense EM data. Sketch illustrates neurite segments obtained from SegEM (Berning et al., 2015), their neighborhood graph (dots and lines), the classification results for connected neurite pieces and synaptic interfaces, and a cell type classifier used to determine the neurite and/or glia type. For details see Methods. (g) Classification result of the neurite/glia type classifiers; red: spine head segments. (h) Reconstruction of all neurons with a cell body and dendrites in the dataset (n=89, total path length of 0.069 m, thus only 2.6% of the path length in the tissue volume, see j). (i) 6 spiny example neurons (SpNs, top, middle) and 2 interneurons (INs, bottom); see Supplementary Material 1 for gallery of all neurons. (j) Quantification of circuit components in the dense reconstruction. Note the majority of circuit path length (total: 2.68 m) is contributed by non-proximal axons (1.78 m, 66.4%), spine necks (0.55 m, 20.5%), and dendritic shafts (0.28 m, 10.4%) not connected to any cell body in the volume. (k) Display of all reconstructed 34,221 axons contained in the dataset. (l) Zoom into the dataset illustrating density of axons (Ax, top, blue-green) and comparison to effective prevalence of dendrites (Dend, bottom, orange-red) at an example location. (m-o) Focused annotation strategy (FocusEM) for directing human annotation queries (Q, red) to ending locations of the automatically reconstructed axon pieces (m, blue). Human annotators were oriented along the axon’s main axis to trace its continuity in webKnossos using flight mode (n, (Boergens et al., 2017)) yielding flight paths of 5.5 ± 8.8 μm length (21.3 ± 36.1 s per annotation, n= 242,271). To detect and correct neurite mergers after automated outgrowth of neurites, locations of chiasmatic shape (o) were detected, and queries (Q) directed from the exits of the chiasma towards its center to determine correct continuities (see Methods). (p,q) Quantification of circuit size and invested work hours for dense circuit reconstructions so far performed in connectomics, and resulting order-of-magnitude improvement provided by FocusEM compared to previous dense reconstructions (q). Fish o.b.: Zebrafish olfactory bulb (Wanner et al., 2016a; Wanner et al., 2016b); M. retina: Mouse retina IPL (Helmstaedter et al., 2013); Fly larva: mushroom body in larval stage of D. melanogaster (Eichler et al., 2017); M. cortex: (Kasthuri et al., 2015) and this study (magenta). Note that only completed dense reconstructions were included in the comparison. Scale bars in c apply to d; h apply to i.

### Cell body-based neuron reconstruction

First, we reconstructed those neurons, which had their cell bodies in the tissue volume (Fig. 1h,i, Supplementary Material 1, n=125 cell bodies, of these 97 neuronal, of these 89 reconstructed with dendrites in the dataset) using a set of simple growth rules for automatically connecting neurite pieces based on the segment-to-segment neighborhood graph and the connectivity and neurite type classifiers (Fig. 1f, see Methods). As a result, we obtained fully automated reconstructions of the neuron’s soma and dendritic processes. Notably with a minimal additional manual correction investment of 9.7 hours for 89 cells (54.5 mm dendritic and 2.1 mm axonal path length), the dendritic shafts of these neurons could be reconstructed without merge errors, but 37 remaining split errors, at 87.3% dendritic length recall (Fig. 1h,i, Supplementary Material 1, see Methods). This reconstruction efficiency compares favorably to recent reports of automated segmentation of neurons in 3D EM data from the bird brain obtained at about 2-fold higher imaging resolution (Januszewski et al., 2018), which reports soma-based neuron reconstruction at an error rate of beyond 100 errors per 66 mm dendritic shafts at lower (68%) dendritic length recall, with a similar resource investment (see Methods).

In addition to the dendritic shafts, the dendritic spines constitute a major fraction of the dendritic path length in cortical neuropil (Fig. 1j). Using our spine head classifier, we found 415,797 spine heads in the tissue volume i. e. density of 0.784 per μm^3^(0.98 per μm^3^ of neuropil, when excluding somata and blood vessels). In order to connect these to the corresponding dendritic shafts we trained a spine neck continuity algorithm that was able to automatically attach 58.9% of these spines (evaluated in the center of the dataset, at least 10 μm from the dataset border) yielding a dendritic spine density of 0.672 per μm dendritic shaft length (comparable to spine densities in the bird brain, (Kornfeld et al., 2017)). However in mammals, the density of spines along dendrites is even higher (about 1 μm'^1^). The remaining spine heads were then attached to their dendritic shafts by seeding manual reconstructions at the spine heads and asking annotators to continue along the spine necks to the dendritic shafts. This consumed an additional 900 hours of human work for the attachment of 98,221 spines, resulting in a final spine density of 0.959 per μm dendritic shaft length.

### Dense tissue reconstruction

The reconstruction of neurons starting from their cell bodies was however not the main challenge. Rather, the remaining processes, that is axons and dendrites not connected to a cell body within the dataset and densely packed in the tissue, constitute about 97% of the total neuronal path length in this volume of cortex (Fig. 1j). To reconstruct this vast majority of neurites (Fig. 1k,l), we first used our connectivity and neurite type classifiers to combine neurite pieces into larger dendritic and axonal agglomerates (see Methods). Then, we took those agglomerates that had a length of at least 5 μm (n=74,074 axon agglomerates), detected their endings that were not at the dataset border and directed focused human annotation to these endings (”queries”, Fig. 1m,n). For human annotation, we used an egocentric directed 3D image data view (”flight mode” in webKnossos), which we had previously found to provide maximized human reconstruction speed along axons and dendrites in cortex (Boergens et al., 2017). Here, however, instead of asking human annotators to reconstruct entire dendrites or axons, we only queried their judgement at the endings of automatically reconstructed neurite parts. To make these queries efficient, we made three additions to webKnossos: We oriented the user along the estimated direction of the neurite at its ending, reducing the time the user needs to orient within the 3D brain tissue; we dynamically stopped the user’s flight along the axon or dendrite whenever another of the already reconstructed neurite agglomerates had been reached; and we pre-loaded the next query while the user was annotating (Fig. 1m,n). With this, the average user interaction time was 21.3 ± 36.1 s per query, corresponding to an average of 5.5 ± 8.8 μm traveled per query. In total, 242,271 axon ending queries consumed 1,978 paid out work hours (i.e. including all overheads, 29.4 s per query).

However, we had to account for a second kind of reconstruction error, so-called mergers, which can originate from the original segmentation, the agglomeration procedure, or erroneous flight paths from human queries (Fig. 1o). In order to detect such mergers, we started with the notion that most of these merger locations will yield a peculiar geometrical arrangement of a 4-fold neurite intersection once all neurite breaks have been corrected (”chiasma”, Fig. 1o). Since such chiasmatic configurations occur rarely in branching neurites, we directed human focused annotation to these locations. First, we automatically detected these chiasmatic locations using a simple heuristic to detect locations at which axon-centered spheres intersected more than three times with the axon (Fig. 1o, n=55,161 chiasmata; for approaches to detect such locations by machine learning, see (Rolnick and Shavit, 2017; Zung et al., 2017)). Then, we positioned the user queries at a certain distance from the chiasma location, pointing inward (Fig. 1o) and then used a set of case distinctions to query a given chiasma until its configuration had been resolved (see Methods for details). Chiasma annotation consumed an additional 1,132 work hours (note that the detection of endings and chiasmata was iterated 8 times for axons, see Methods, and that in a final step we also detected and queried 3-fold neurite configurations to remove remaining mergers). With this we obtained in summary a reconstruction of 2.72 meters of neuronal processes (Fig. 1p, 0.89 meters of dendrites (including 0.55 meters of spine necks) and 1.76 meters of axons) with a total investment of 3,981 human work hours-about 10 times faster than a recent dense reconstruction in the fly larval brain ((Eichler et al., 2017), Fig. 1q), about 20 times faster than the previous dense reconstruction from the mammalian retina ((Helmstaedter et al., 2013)), and about 25 times faster than the previous dense reconstruction from mammalian cortex (Kasthuri et al., 2015).

We then measured the remaining reconstruction error rates in this dense neuropil reconstruction. Since the following of neurites in dense neuropil is much more difficult than the reconstruction of dendrites and proximal axons from the cell body (Fig. 1h,i) we expected this error rate to be substantially higher. In fact, when we quantified the remaining errors in a set of 10 randomly chosen axons we found 12.8 errors per millimeter of path length (of these 8.7 per millimeter continuity errors, see Methods). This is indistinguishable from the error rates previously found in fast human annotations (Boergens et al., 2017; Helmstaedter et al., 2011; Helmstaedter et al., 2013).

### Connectome Reconstruction

Given the reconstructed pre-and postsynaptic neurites in the tissue volume, we then went on to extract their connectome. For this we used SynEM (Staffler et al., 2017) to detect synapses between the axonal presynaptic processes and the postsynaptic neurites (for non-spine synapses, improvements to SynEM were made to enhance precision and recall, see Methods). Since we were interested in analyzing the subcellular specificity of neuronal innervation, we had to also classify which of the post-synaptic membranes belong to cell bodies; to classify spiny dendrites as belonging to excitatory cells, smooth dendrites belonging to interneurons; and to detect axon initial segments and those dendrites that were likely apical dendrites of neurons located in deeper cortical layers. We developed semi-automated heuristics to detect these subcellular compartments (Fig. 2a-d, see Methods for details).

**Fig.2.**
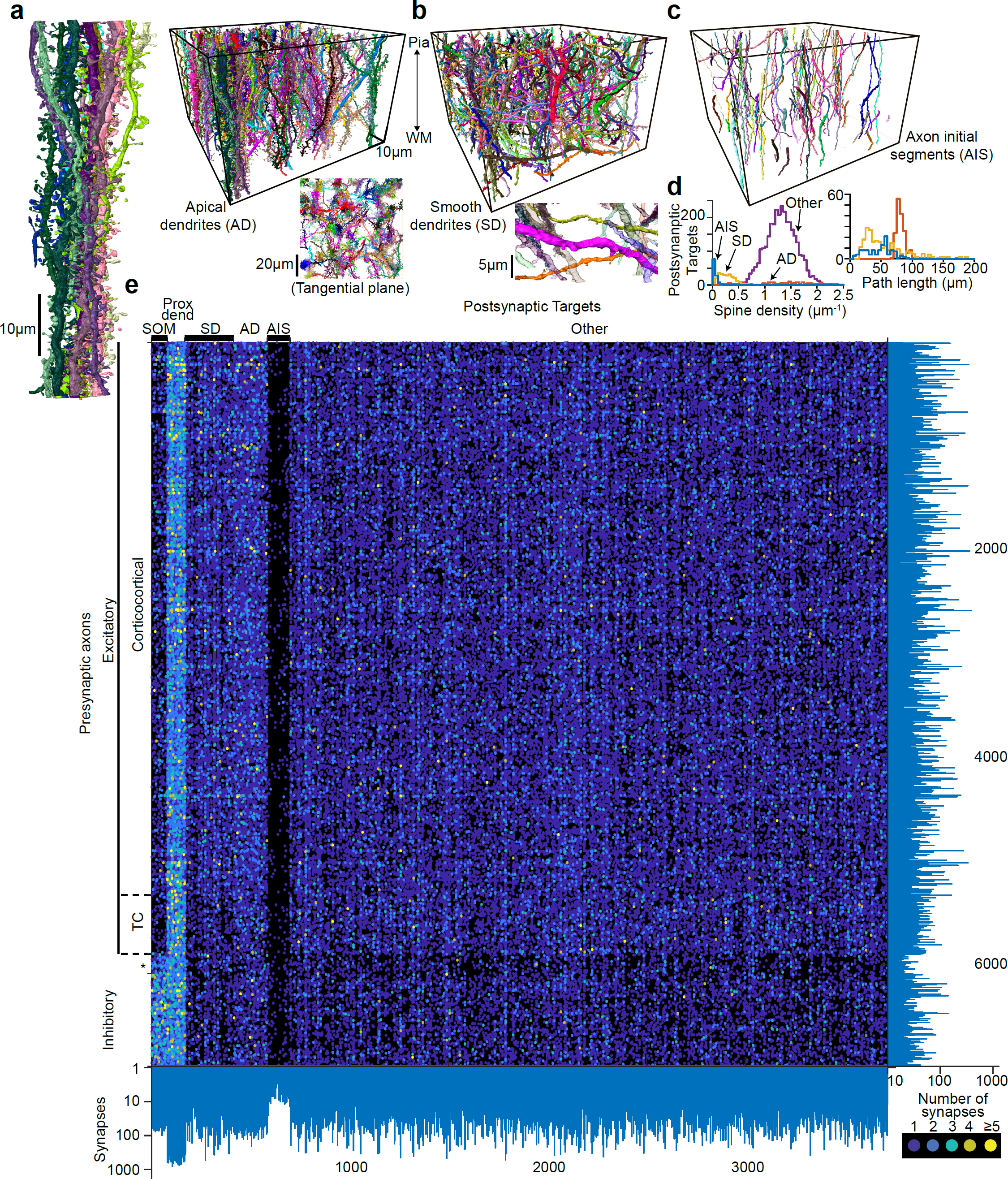
Postsynaptic target classes and dense cortical connectome. (a-d) Display of all apical dendrites (AD, a, magnified one apical dendrite bundle (left), and top view in tangential plane illustrating AD bundles), smooth dendrites (SD, b, magnification inset illustrates low rate of spines), axon initial segments (AIS, c) and their respective path length and spine density distributions (d). **(e)** Display of connectome between all axons (n=6,979) and postsynaptic targets (n=3,719) in the volume with at least 10 synapses, each; total of 153,171 synapses (of 388,554 synapses detected in the volume). Definition of postsynaptic target classes, see (a-d); definition of presynaptic axon classes: see Fig. 3. Note that for AIS also those with less than 10 input synapses are shown. SOM: neuronal somata; Prox. Dend.: proximal dendrites connected to a soma in the dataset; asterisk, remaining unassigned axons (see Fig. 2b).

With this we obtained a connectome between 34,221 pre-synaptic axonal processes and a total of 11,400 post-synaptic processes (i. e. somata, axon initial segments and dendrites). Restricting the connectome to those pre-and postsynaptic neurites that established at least 10 synapses each yielded a 6,979 by 3,719 connectivity matrix, reporting the number of synapses established between each pair of pre-and postsynaptic neurites (Fig. 2e). The postsynaptic processes comprised 80 somata, 246 smooth dendrites, 169 apical dendrites, and 116 axon initial segments (Fig. 2e, for AIS also those with less than 10 input synapses are shown). The dendrites of soma-based neuron reconstructions were labeled as proximal dendrites. In addition, we automatically determined for each synapse whether it was established onto a spine head, dendritic shaft or soma.

### Synaptic specificity

Then we investigated whether based solely on connectomic information (Fig. 2) we could extract the rules of subcellular innervation specificity described for inhibitory axons in the mammalian cortex (for a review see (Kubota et al., 2016)), and whether such synaptic specificity could also be found for excitatory axons. We first measured the preference of each axon for innervating dendritic spine heads versus dendritic shafts and other targets (Fig. 3a,b). In the mammalian brain, most axons of inhibitory interneurons preferentially innervate the dendrites’ shafts or neuronal somata (see e.g. (Kubota et al., 2016)), and most excitatory glutamatergic axons preferentially innervate the spine heads of dendrites (Feldmeyer et al., 2002; Shepherd and Harris, 1998). Accordingly, in our dense data, we found that the fraction of primary spine synapses per axon (out of all synapses of that axon) has a clear peak at about 80% (Fig. 3a,b), allowing the identification of spine-preferring, likely excitatory axons with at least 50% primary spine innervations. Similarly, we identified shaft-preferring, likely inhibitory axons with less than 20% primary spine innervations. Together this yielded 6,449 axons with clear shaft or spine preferences. For the remaining n=528 axons with primary spine innervations above 20% and below 50%, we first wanted to exclude remaining mergers between excitatory and inhibitory axons (that would yield intermediate spine innervation rates) and split these axons at possible merger locations (at least 3-fold intersections). Of these, 338 now had at least 10 synapses and spine innervation rates below 20% or above 50%. The remaining n=192 axons (2.75% of all axons with at least 10 synapses) were not included in the following analyses. This together yielded n=5,894 excitatory and n=893 inhibitory axons in our data.

**Fig.3.**
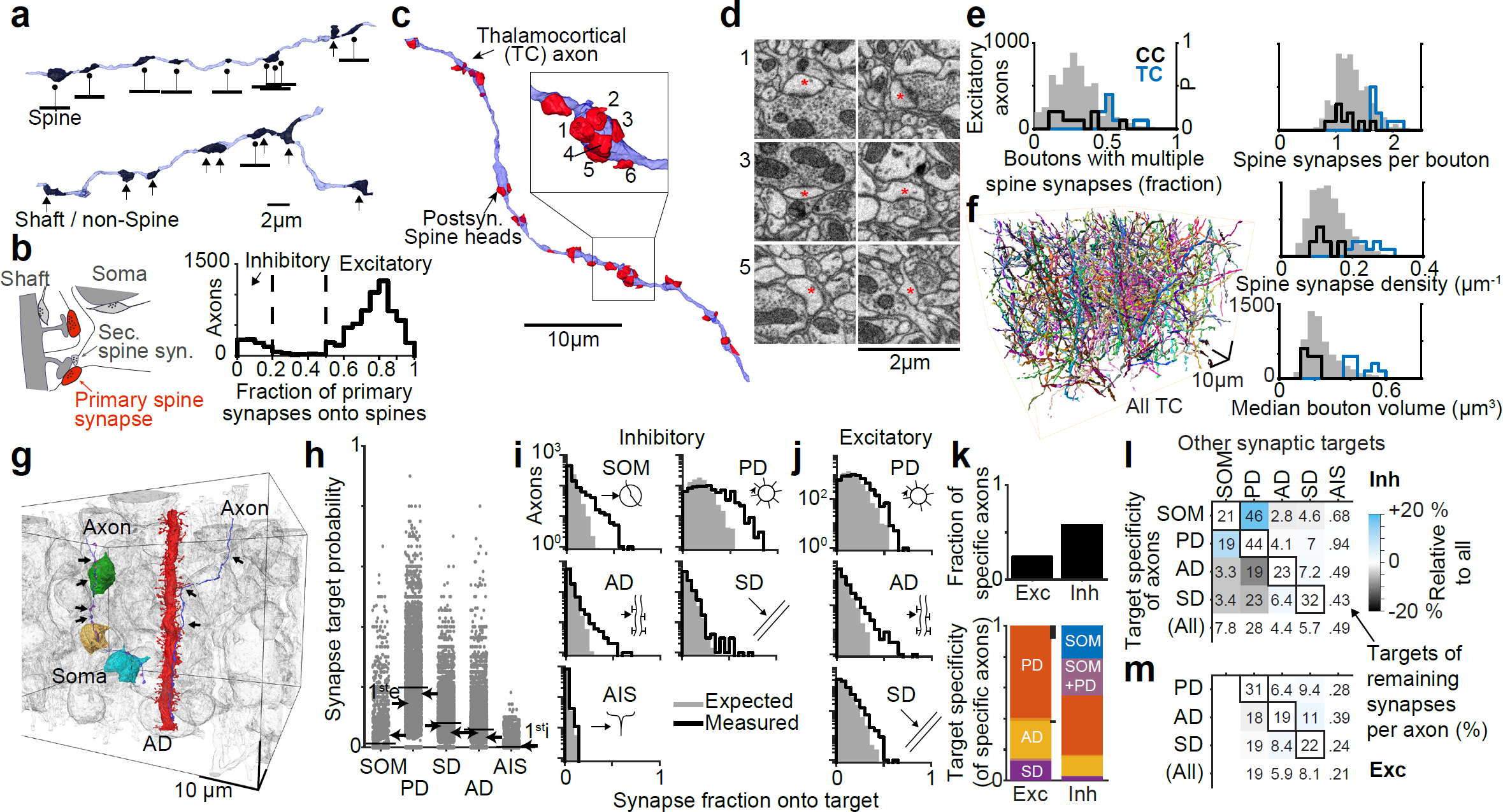
Connectomic definition of axon classes. (a) Example axons with high (top) and low (bottom) fraction of output synapses made onto dendritic spines. (b) Distribution of spine targeting fraction over all n=6,979 axons; dashed lines indicates thresholds applied to distinguish non-spine preferring, likely inhibitory (i, <20% spine innervation, n=893, 12.8% of all axons) from spine-preferring, mostly excitatory (e, >50% spine innervation, n=5,894, 84.5% of axons) axons. Sketch illustrates measurement of spine fraction as fraction of primary spine innervations out of all other synapses. (c-e) Identification of thalamocortical axons by previously established criteria (Bopp et al., 2017) relating to their high frequency of multiple-target large boutons (example in c,d; red asterisks, postsynaptic spine heads of same TC bouton). Quantification of these properties for all excitatory axons (gray shaded) and test sets of clear thalamocortical (TC, blue) and cortico-cortical (CC, black) axons (relative distributions, right axes). (f) Resulting set of n=569 TC axons in dataset. (g-m) Connectomic evidence for subcellular target specificity of certain axon classes. (g) Two example axons innervating three somata (SOM, left, n=8 synapses onto somata of 18 total) and an apical dendrite (AD, right, n=2 synapses onto AD of 13 total), respectively. All other cell bodies and ADs in gray. (h) Overall fraction of synapses onto SOM, PDs, ADs, SDs, AIS for all n=6,979 axons. Arrows indicate binomial probabilities over axons to establish at least one synapse onto the respective target (1^st^e for excitatory axons, arrow pointing right; 1^st^i for inhibitory axons, arrows pointing left). Black lines, average over axons. (i) Comparison of predicted synapse fraction onto target classes per inhibitory axon based on the binomial probability (p1sti, see h) to innervate the target at least once (gray shaded) and measured distribution of synapse fractions onto targets (black lines). Note longer tail of measured innervation fraction in measured ***VS.*** expected data for SOM, PD, AD and SD, but not AIS. (j) Same analysis as in (i) for excitatory axons indicates AD, PD-and SD-specificity. (k) Fraction of target-specific excitatory (Exc) and inhibitory (Inh) axons identified using the false detection rate criterion (q=20%, (Storey and Tibshirani, 2003)). Black bars indicate TC axons. Mixed colors indicate axons specific for both SOM and PD. (l,m) 2^nd^ order innervation pattern for target-specific axons; numbers report fractional innervation by non-class specific, remaining synapses per axon; colors indicate under-(blue) or over-(red)-frequent innervation. Diagonal entries report fraction of synapses onto same specific target (black boxes). The conditional dependence of target innervation especially for the inhibitory axon classes in this 2^nd^ order analysis serves as *post-hoc* evidence of the connectomic definition of axonal types as shown in (a-k).

Previous reports have described that a subset of excitatory axons in cortical L4 preferentially target the shafts of dendrites in some species (Lubke et al., 2000; McGuire et al., 1984): a study of L4 spiny neurons’ axons in juvenile rat found preferential innervation of small-caliber dendritic shafts (and only 27% of synapses onto spines, (Lubke et al., 2000)), and in cat visual cortex, a subset of corticortical excitatory axons from layer 6 has been described to establish preferentially shaft synapses onto spiny dendrites in L4 at the end of short axonal branches, yielding boutons terminaux (Ahmed et al., 1994; McGuire et al., 1984)). To check whether these axons would confound our assignment of shaft-preferring axons as inhibitory, we randomly selected 20 shaft synapses onto spiny dendrites and manually reconstructed the presynaptic axons with their output synapses. We first asked whether any of these axons would preferentially establish boutons terminaux onto shafts, as described in cat, but found no such axon, indicating that this innervation phenotype comprises less than 5% or is absent in our data from mouse L4 (compare to the estimate of more than 40% of such inputs in cat L4 (Ahmed et al., 1994)). We then checked whether any of the 20 axons showed both a preference for shaft innervation and in a minority of cases any clear primary spine head innervation, as described for the L4 axons in juvenile rat. We found no such example, indicating that none of the shaft-preferring axons was excitatory. This is consistent with data from cat which suggested that L4 spiny axons preferentially target spines (Ahmed et al., 1994). Together, we conclude that in our data from mouse L4, excitatory axons preferentially establish primary spine head innervations (Fig. 3b) and inhibitory axons preferentially innervate the shafts of dendrites.

Within cortical layer 4, the two main types of excitatory synaptic input are afferents from the thalamus (thalamocortical inputs, TC) and intracortical inputs (corticocortical, CC). In order to distinguish between corticocortical and thalamocortical excitatory axons (Fig. 3c-f), we used previously established criteria about the frequency of multi-target boutons, bouton size and the number of targets per bouton that had been shown to identify TC inputs in layer 4 of mouse S1 cortex ((Bopp et al., 2017); Fig. 3c-f, see Methods). Using these, we extracted the likely thalamocortical (TC) axons (Fig. 3e,f; n = 569, 9.7% of excitatory axons).

We then determined for each of the subcellular synaptic target classes (somata (SOM), axon initial segments (AIS), apical dendrites (AD), smooth dendrites (SD), proximal dendrites (PD), see Fig. 2 and Fig. 3g) the per-synapse innervation probability that would best explain whether an inhibitory axon establishes at least one synapse onto each of these targets (these inhibitory “first-hit” binomial innervation probabilities were 4.2% (SOM), 17.8% (PD), 4.9% (SD), 3.3% (AD), and 0.5% (AIS), see Methods, Fig. 3h). We then computed the expected distribution of synapses per axon made onto each target class assuming the second-hit, third-hit, etc. innervation probabilities are the same as the probability to establish at least one synapse onto that target. When comparing these target distributions to the actually measured distributions of synapses per axon onto each target class (Fig. 3i), we found that inhibitory axons established enhanced specificity for cell bodies (p=2.4x10^-34^, n=893, one-sided Kolmogorov-Smirnov test), proximal dendrites (p=6.0x10^-77^), apical dendrites (p=2.5x10^-4^) and smooth dendrites (p=1.7x10^-3^), but no enhanced specificity for axon initial segments in L4 (p=0.648, note that AIS are synaptically innervated by 0.172 input synapses per μm AIS length; but these innervations are not made by certain axons specifically, unlike in supragranular layers (Taniguchi et al., 2013)).

When performing the same analysis for excitatory axons (Fig. 3j), we found clear target specificity for apical dendrites (p=2.5x10^-34^, Fig. 3j), for smooth dendrites (p=7.6x10^-25^) and for proximal dendrites (p=1.3x10^-169^). Thalamocortical axons, to the contrary, show indication of target specificity for proximal dendrites (p=2.5x10), but not for apical (p=0.019) or smooth dendrites (p=0.723).

These results provide statistical connectomic evidence for the existence of target-specific wiring of inhibitory and excitatory axons in cortical layer 4. Next, we wanted to determine the fraction of inhibitory and excitatory axons that had an unexpectedly high synaptic preference for one (or multiple) of the subcellular target classes. For this, we determined for each axon the probability that its particular synaptic target choices had originated from a simple chance drawing given the first-hit probabilities (Fig. 3h), or whether it showed additional specificity. Here, we used the false detection rate criterion used for the determination of significantly expressed genes (q value, (Storey and Tibshirani, 2003), see Methods). As a result, we obtained lower bounds on the fractions of axons in the tissue that are specifically innervating the various subcellular target classes (Fig. 3k; 58.0% of inhibitory and 24.4% of excitatory axons). Of those inhibitory axons found to be target-specific, about 83% of axons are specific for somata or proximal dendrites, about 14% for apical dendrites, and about 3% for the smooth dendrites of other interneurons. Interestingly, we also found subsets of excitatory axons with subcellular target specificity: of those excitatory axons with significant synaptic target specificity, about 28% are specific for apical dendrites and about 14% for smooth dendrites. Furthermore, at least 24.7% of thalamocortical axons specifically innervated proximal dendrites (Fig. 3k).

Finally, we asked whether the specificity of axons towards one particular synaptic target yields an enhanced (or suppressed) innervation of other synaptic targets (i.e., whether axons exhibit conditional, higher-order synaptic specificity, Fig. 3l,m). For example, given axons that show enhanced innervation of somata, would the target distribution of the non-somatic synapses of these axons be random, or would these remaining synapses show additional target preferences or target suppression (Fig. 3l,m)? For this, we analyzed all axons of certain target specificity as identified before (Fig. 3k), excluded synapses of these axons onto their specifically innervated target, measured the fractions of remaining synapses onto the other target classes and compared them to the average innervation rate over all axons (Fig. 3l,m). We found that inhibitory axon subpopulations with soma-and proximal dendrite-specificity are overlapping, and axons with specificity towards apical and smooth dendrites exhibit suppressed innervation of the proximal dendritic/somatic targets and vice versa (Fig. 3l). Excitatory axons show only very weak conditional innervation preference (Fig. 3m).

Together these results represent the patterns of subcellular synaptic innervation rules exhibited in a local cortical circuit in layer 4. It should be noted that the definition of presynaptic axonal types was performed relying only on connectomic data; not on expression markers or cell morphology. Such a cell type classification based on local connectomic data alone has been successful in the mammalian retina before (connectomic definition of a third subtype of type 5 bipolar cells (Helmstaedter et al., 2013)) – but whether it would be possible in dense mammalian cortical data was not clear *a-priori.* The fact that the connectomically defined axonal classes exhibit additional higher-order innervation preference (Fig. 3l) further indicates that these are in fact valid axonal type definitions.

### Geometric explainability of synaptic innervations

We were now able to ask whether these local connectivity rules (Fig. 3) could have been derived solely from the geometry of axons and dendrites. The question to what degree the trajectories of axons and dendrites are already predictive of synaptic innervation in the cerebral cortex has been controversially debated (Binzegger et al., 2004; Braitenberg and Schüz, 1998; Kasthuri et al., 2015; Lee et al., 2016; Markram et al., 2015), and an assumption of random innervation has been put forward (Peters’rule, (Braitenberg and Schüz, 1998)) and used for massive simulation initiatives (Markram et al., 2015). While connectomic examples of non-random innervation in the cortex are documented (da Costa and Martin, 2009; Kasthuri et al., 2015; Mishchenko et al., 2010; Schmidt et al., 2017), a rigorous analysis within dense cortical neuropil of sufficient scale is missing.

We first compared the fraction of synapses made onto the subcellular target classes with the fraction of membrane surface attributed to these subcellular domains, sampled in cubes of ~5 μm edge length within the dataset volume (Fig. 4a). The membrane surface fraction deviated up to a factor of 2 from the actual synapse fraction, both underestimating (apical dendrites, Fig. 4a) and overestimating (somata) the synaptic innervation.

**Fig.4.**
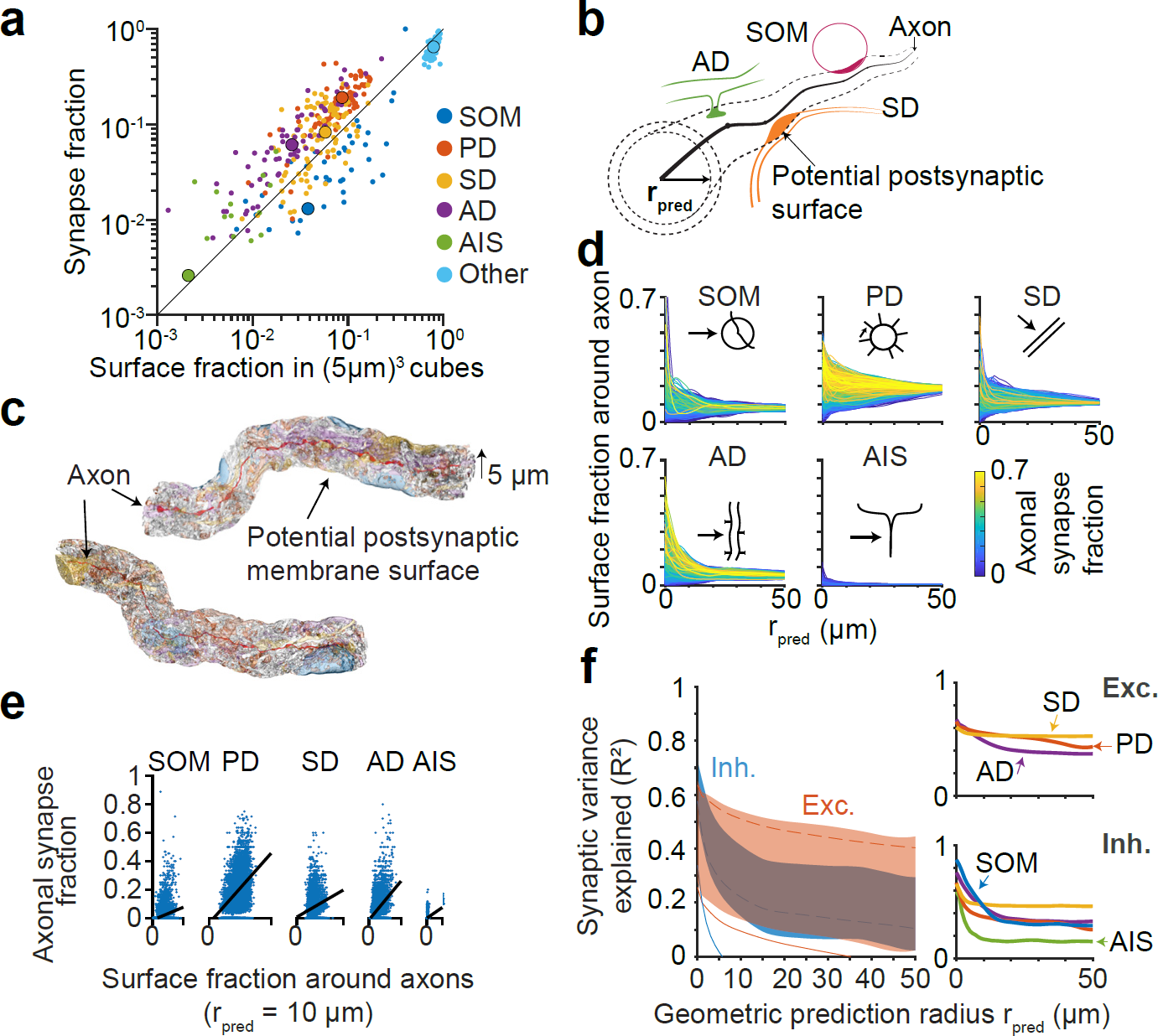
Contribution of neurite geometry and postsynaptic membrane availability to cortical wiring. (a) Comparison between the fraction of membrane surface area attributed to one of the subcellular target classes (colors, see Figs. 2,3) and the actual fraction of synapses made onto these target classes, sampled in cubes of ~5 μm edge length (dots) distributed across the dataset volume (average over entire dataset, large circles). **(b)** Sketch of the axon-based measurement of the fraction of surface area of the various subcellular target classes (colors) within a distance r_pred_*f*from a given axon (black). **(c)** Example surfaces around two axons (Ax1, Ax2, same as in Fig. 3g) with r_pre_*d* = 5 μm (shaded colors: target classes as in a). Ax_1_ showed soma, Ax_2_ AD specificity in the analyses in Fig. 3. **(d)** Corresponding surface fraction of somata and apical dendrites for the two axons in c, in dependence of r_pred_. **(e)** Same as (d) for all n=6,979 axons in the dataset, shown separately for the target classes (see symbols). Colors denote the fraction of synapses of a given axon that innervate the respective target class. **(f)** Relation between the surface fraction around all axons and the synaptic innervation by these axons for each target class, shown for r_pred_ of 10 μm. Black lines show linear regression to obtain optimal model parameters for geometrical innervation prediction in g. **(g)** Coefficient of determination (R^2^) reporting the fraction of synaptic innervation variance (over all axons, see f) explained by an innervation model using the available postsynaptic surface area around axons (shaded area; red, excitatory axons; blue, inhibitory axons; lower end of shades indicates prediction; upper ends indicate correction by the variance contributed by the multinomial sampling of targets along axons, see Methods). Insets (right) show sampling-corrected predictive power of excitatory (top) and inhibitory (bottom) axons for the innervation of target classes. This analysis refutes a random geometry-based innervation rule for axons in dense cortical neuropil.

We then investigated whether the postsynaptic membrane surface available within a certain radius r_pre_*d* around a given axon (Fig. 4b,c) would be a better predictor of synaptic innervation for that given axon. For this we measured the available membrane surface belonging to the 5 subcellular target classes around all axons (Fig. 4d).

We then used a logistic multinomial regression model to predict synaptic innervation from the availability of membrane surface attributed to the target classes around the 6,979 axons (Fig. 4e). In this we assumed that the precise axonal trajectories were known (corresponding to perfect alignment of axonal and dendritic reconstructions), and that the number of synapses per axon was given. Based on this, we computed the coefficient of determination (R^2^) reporting the fraction of axonal synaptic innervation variance that could be explained purely based on the geometrical information (Fig. 4f). For this we subtracted the variance originating from the multinomial sampling of a concrete innervation target per synapse and axon (see Methods), thus reducing the variance that has to be explained by the geometric model. Yet, even using these favorable conditions, at a prediction radius of 10 μm, only up to 41% of inhibitory innervation and about 54% of excitatory innervation variance was accounted for by the geometric model (Fig. 4f). This lack of geometrical predictability was present for all types of axons (Fig. 4f). Notably, commonly used integration scales for geometrical connectomic prediction (25 μm, (Binzegger et al., 2004; Lubke et al., 2003; Meyer et al., 2010; Oberlaender et al., 2012a)) provided only about 34% explained variance of actual synaptic innervation for inhibitory and only about 50% for excitatory axons under the rather optimal predictive conditions as described above.

### Dendritic and axonal synapse positioning

We next investigated the distribution of input and output synapses along the soma, dendrites and axons of the excitatory L4 neurons (Fig. 5). Up to about 20 μm from the soma, almost no spines are established (Fig. 5a). While the total number of excitatory synapses increases substantially until about 50 μm from the soma, the fraction of excitatory input contributed by TC axons stays remarkably constant at about 12% from 20 μm onwards (Fig. 5b). The inhibitory-excitatory synaptic input ratio (i/(i+e)) drops from almost 100% to about 15% within 50 μm, and further decreases to about 7% at 100 μm from the soma (Fig. 5b).

**Fig.5.**
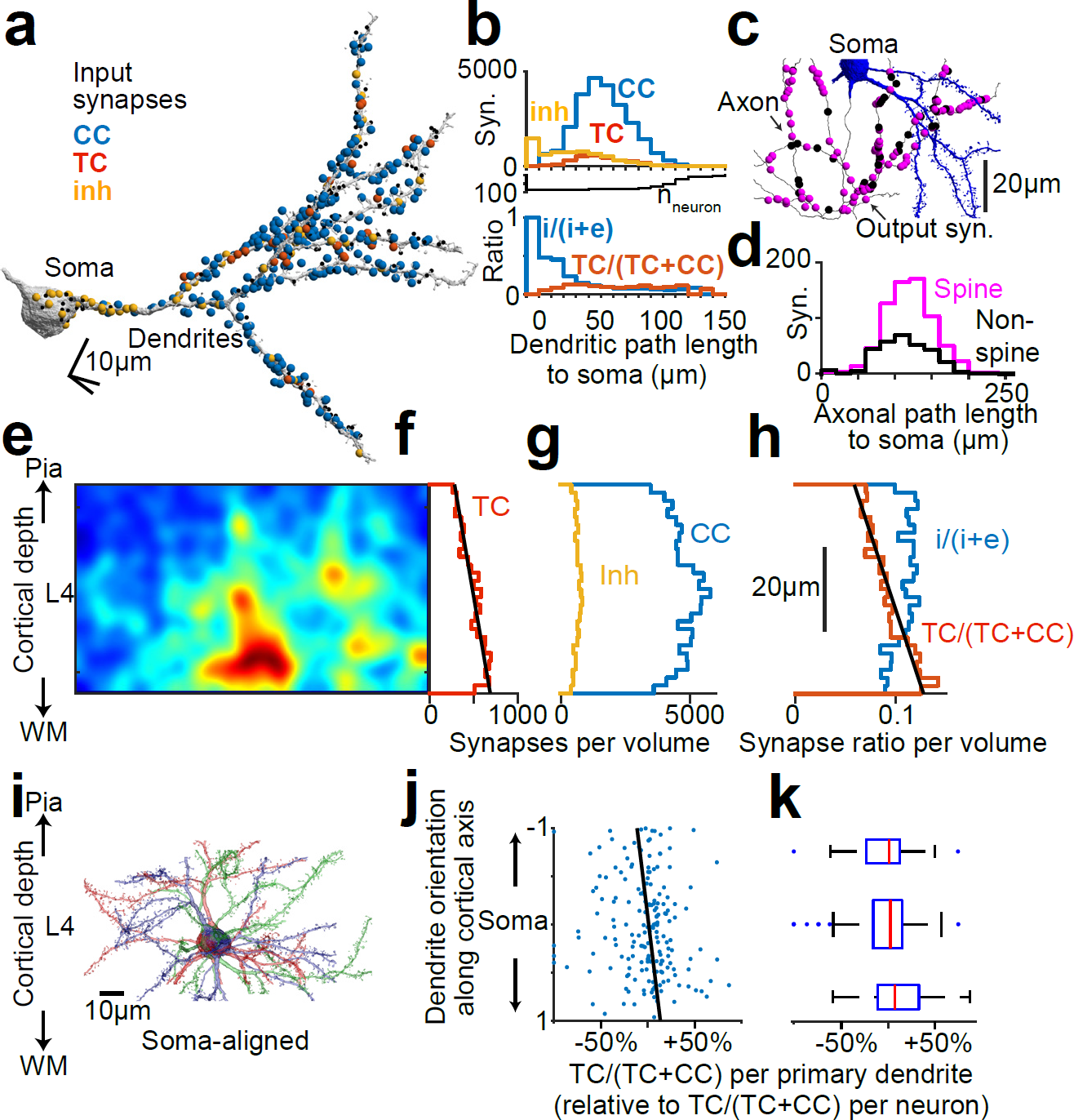
Distribution of synapses along dendrites and axons, and variability of synaptic input composition. (a) Neuron with cell body and primary dendrites; all input synapses are indicated, colored according to the type of presynaptic axon (yellow: inhibitory; red: TC, blue: CC; black, axon with less than 10 synapses). **(b)** Distribution of input synapses (top) and resulting inhibitory/excitatory balance and TC input fraction (bottom) summed over all neurons in the dataset (n=90 neurons, n=183 primary dendrites, total of n=47,552 synapses) as function of dendritic path length to soma. Black line indicates number of neurons (n_neuron_) contributing to the respective distance bin. **(c)** Neuron with soma and local axon collaterals, all output synapses are indicated according to the target (magenta: spine; black: non-spine). **(d)** Average distribution of spine and non-spine targets along path length of L4 output axons refuting path-length dependent axonal synapse sorting (PLASS) for L4 of mouse S1 (compare (Schmidt et al., 2017) for mammalian cortex). **(e-h)** Distribution of TC synapses within L4 dataset (e) shows gradient along the cortical axis (f), which is absent for inhibitory or CC synapses (g). **(h)** Resulting gradient in TC synapse fraction (increase by 83% from 7.0% to 12.8% (+5.8%) within 50 μm along the cortical axis; line fit, p<1.1x10, n=134,537 synapses). **(i-k)** Analysis of the variability of TC input onto the primary dendrites of neurons possibly resulting from the TC synapse gradient (h): example reconstructions (i) aligned to the somata; (j) fraction of excitatory input synapses originating from TC axons evaluated for each primary dendrite, plotted according to the direction of the dendrite in relation to the cortical axis (-1: dendrite aligned towards the pia; +1: dendrite aligned towards the white matter). The TC input fraction (TC/(TC+CC)) of each dendrite was compared to the TC input fraction of its entire parent neuron, ratios are shown. (k) Summary analysis of relation between dendrite direction and relative TC input fraction. Note that at the level of single dendrites, TC input fraction is determined by the dendrites’ orientation relative to the cortex axis (k, 1.28-fold higher relative TC fraction for downwards pointing dendrites (projection >0.5) than upwards pointing dendrites (projection <-0.5), n=183, p=0.026, two-sided t-test for dendrites with a normalized absolute projection >0.5; bars correspond to dendrites from projection ranges-1.0.5;-0.5.0.5; 0.5.1, respectively). This data indicates a synaptic input mixing that enhances synaptic input variability at the level of single dendrites.

To study the positioning of synapses along excitatory axons, we used those axons leaving the cell bodies to ask whether synapses were sorted along the axonal path according to their target (Fig. 5c,d). While target-sorted placement of output synapses along axons had been theoretically predicted and found in non-mammalian species (Carr and Konishi, 1988, 1990; Jeffress, 1948; Kornfeld et al., 2017), the discovery of sorted synaptic arrangements along axons in layer 2 of the mammalian medial entorhinal cortex was a surprise (Schmidt et al., 2017). Here, we found no evidence for path-length dependent axonal synapse sorting (PLASS) in layer 4 of mouse somatosensory cortex (Fig. 5d), excluding PLASS as a ubiquitous cortical wiring principle, but leaving open the possibility that it could be a feature of nongranular cortex.

### Synaptic input variability

Functional recordings of cortical neurons *in-vivo* show a remarkable variability of responses between neurons but also between stimulus exposures for a given neuron (Brecht and Sakmann, 2002; Kerr et al., 2007; Kerr et al., 2005; Ohki et al., 2005; Stosiek et al., 2003). A heterogeneous sampling of available synaptic inputs at the level of neurons and dendrites equipped with non-linear functional properties (Lavzin et al., 2012) could be one mechanism generating such variable functional responses. Using our dense synaptic input data, we wanted to next analyze the variability of synaptic input composition in L4 neurons (Fig. 5e-n).

We first noticed that the density of thalamocortical synapses had a substantial dependence on cortex depth (Fig. 5e-h): the absolute density of TC synapses in the volume increased by about 93% over 50 μm cortex depth (Fig. 5e,f; the TC excitatory synapse fraction TC/(TC+CC) increased by 82.6%, corresponding to an absolute increase in the TC synapse fraction of 5.8% per 50 μm cortex depth, Fig. 5h). This gradient is consistent with light-microscopic analyses of TC synapses showing a decrease of TC synapse density from lower to upper L4 (Garcia-Marin et al., 2013; Oberlaender et al., 2012b; Wimmer et al., 2010) which is most substantial when analyzed at the level of single VPM axons (Oberlaender et al., 2012b). Neither the inhibitory nor the corticocortical synapse densities showed a comparable spatial profile (Fig. 5g).

We wanted to understand how the synaptic TC gradient is mapped onto the input of L4 neurons along the cortex axis (Fig. 5i-k). One possibility was that the TC synapse gradient (Fig 5e,f,h) is used to enhance the variability of synaptic input composition between different primary dendrites of the L4 neurons such that a neuron’s dendrites pointing upwards towards the pia would sample relatively less TC input than dendrites pointing towards the white matter. Alternatively, synaptic specificity mechanisms (as in Fig. 3) could be used to counterbalance this synaptic gradient and equilibrate the synaptic input fractions on the differently oriented dendrites. Our analysis (Fig. 5j,k) shows that in fact, even for single primary dendrites, TC input fractions vary 1.28-fold between dendrites pointing upwards towards the cortical surface vs downwards towards the white matter (TC input fractions of each dendrite were corrected for the entire neuron’s TC input fraction for this analysis, see Methods). This finding of per-dendrite input variation points to a circuit configuration in which TC input variability is enhanced between and within neurons of the same excitatory type in cortical layer 4.

The finding of a substantial TC synapse gradient along the cortical axis within L4 is interesting, since the fractional thalamocortical innervation of spiny neurons in L4 has been a matter of extensive scientific investigation (Ahmed et al., 1994; Benshalom and White, 1986; Bopp et al., 2017; da Costa and Martin, 2009; Garcia-Marin et al., 2017; Latawiec et al., 2000; White, 1989; White and Hersch, 1981), with results of TC input fraction ranging from less than 10% (da Costa and Martin, 2009) to up to 20% (Benshalom and White, 1986; White, 1989) of synaptic excitatory input contributed from the thalamus in layer 4 of sensory cortex. While most differences have so far been attributed to species differences (Bopp et al., 2017), our data supports the view that cortical depth within layer 4 may be a key determinant of TC input fraction (Garcia-Marin et al., 2017). Our data also emphasizes the heterogeneity of synaptic input for cortical neurons of similar type.

### Connectomic signature of synaptic plasticity

Finally we used the unprecedented magnitude of synaptically coupled axons and dendrites in this dense cortical volume to look for a potential connectomic signature of previous episodes of synaptic plasticity. In one well-established concept of neuronal plasticity, the timing of action potentials in pre-and postsynaptic neurons is the key determinant of synaptic weight change (Hebb, 1949; Markram et al., 1997). Since synaptic strength has been shown to be correlated to the size of synaptic specializations (postsynaptic density, spine head volume, (Harris and Stevens, 1988, 1989) and axon-spine interface (ASI) area (de Vivo et al., 2017)), the analysis of the similarity of synaptic size for pairs of pre-and postsynaptic neurites that establish more than one joint synapse can be used to obtain evidence for previous episodes of synaptic weight assimilation between these joint synapses. In the hippocampus, where activity-dependent synaptic weight increase is consistently found (LTP), this argument has been employed to investigate the storage capacity of synapses (Bartol et al., 2015; Bromer et al., 2018), and joint-synapse size homogeneity has been described for clustered synaptic input (Bloss et al., 2018). Here, we wanted to determine whether such synaptic size homogenization can be quantitatively determined in cortical layer 4, in which only long-term synaptic depression has been found (Egger et al., 1999), and to determine upper and lower bounds on the fraction of the circuit that can have recently undergone such synaptic weight adaptation.

For this, we first searched for all pairs of axons and dendrites that established more than one joint axon-to-spine synapse in the volume (Fig. 6a-c, n=6,602; of these n=6,045 with 2, n=474 with 3, and n=83 with 4 and more synapses; we call them joint synapses in the following; this analysis was carried out both in the original axon reconstruction and in a reconstruction with all axons split at potential branch points and merger points; the latter served as control for the influence of merge errors on the results; in both cases the reported effects were found; the reported numbers are from the control case). We then measured the size of the axon-spine interface (ASI, (de Vivo et al., 2017; Staffler et al., 2017)) for all synapses, those with 2 joint synapses, etc. (Fig. 6d). Surprisingly, mean synaptic size increased by 1.15-fold per additional joint synapse. Given the absence of electrophysiological evidence for LTP in L4 (Egger et al., 1999), this finding could correspond to the subset of extremely strong unitary synaptic connections in L4 reported by (Feldmeyer et al., 1999) which had a unitary synaptic efficacy of up to 10 mV, allowing to elicit postsynaptic APs based on one synaptically connected presynaptic neuron. Alternatively, this data could indicate that in a subset of the L4 circuit, LTP is established. In this case, our data would support a model in which additional synapses are added when previous coincident pre-and postsynaptic activity has already strengthened the existing joint synapses. This could also indicate that the more joint synapses are established between LTP-enabled connections, the more likely it is that the single presynaptic axon elicits APs in the postsynaptic neuron, enhancing possible synapse strength increase-and giving rise to the reported extremely large unitary connections (Feldmeyer et al., 1999).

**Fig.6.**
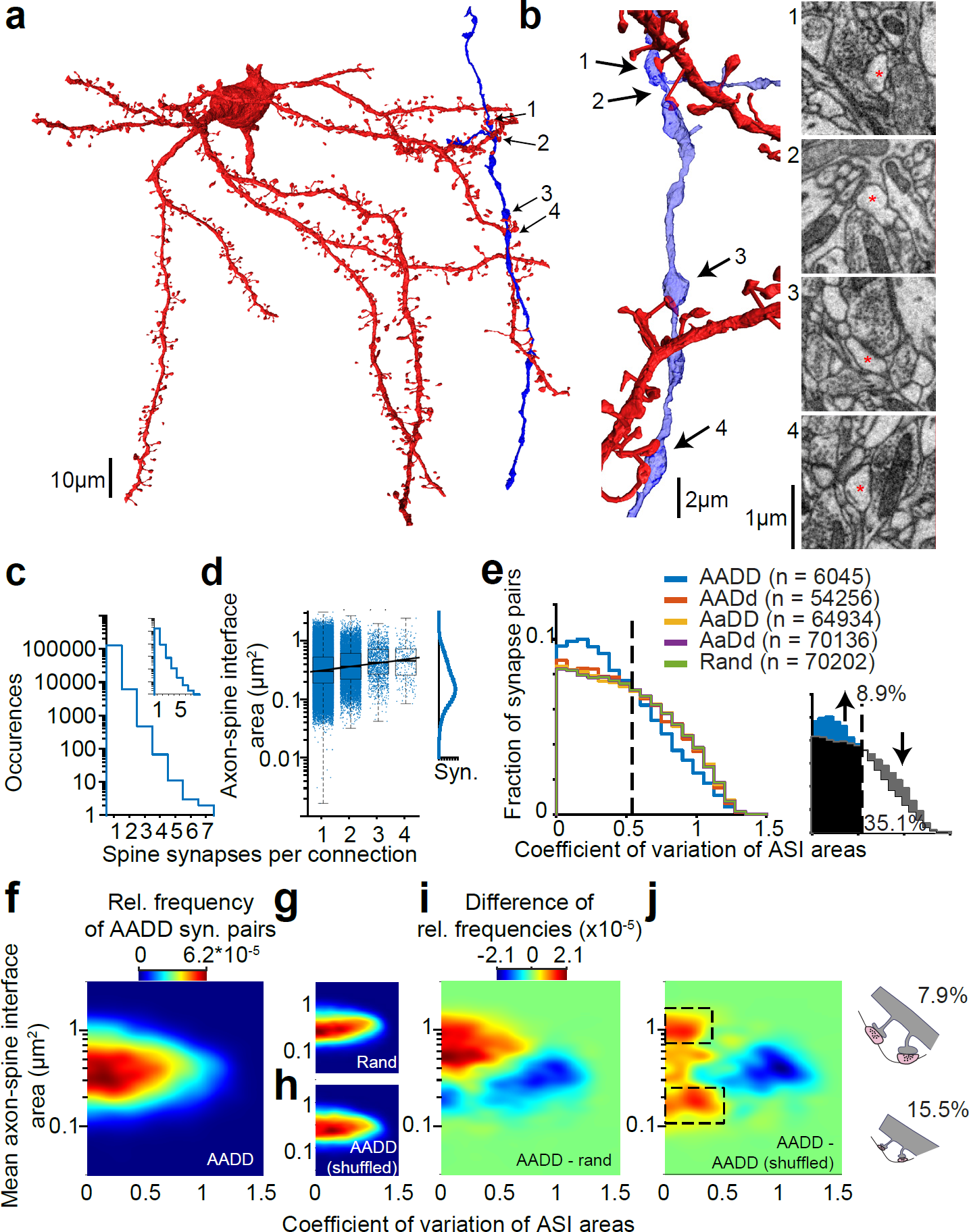
Analysis of multisynaptic connections for spine synapse size homogenization possibly induced by Hebbian learning. (a,b) Example of an axon innervating the dendrites of a postsynaptic neuron 4 times within the dataset (b; red asterisks indicate innervated spine heads). **(c)** Frequency of multi-hit connections. Note that n=6,045 connections involved 2 synapses, and n=557 at least 3; numbers are for control reconstruction set, in which all axons were split at branch points to avoid contamination by axon mergers (unsplit axons in inset); see Methods. **(d)** Distribution of axon-spine interface (ASI, (de Vivo et al., 2017; Staffler et al., 2017)) size over single and multi-hit connections indicating larger synapses in multisynaptically connected pairs. Inset: overall single synapse ASI size distribution following a lognormal distribution. **(e)** Analysis of synapse size similarity for pairs of synapses drawn randomly (rand, green) and from unrelated axons and dendrites (AaDd, purple), pairs of same axon but different dendrites (AADd, red), same dendrite but different axons (AaDD, yellow), and same axon-same dendrites (AADD, blue). The distribution of synaptic size variability (coefficient of variation) for pairs of randomly drawn synapses and for those involved in AADD connections shows that low-variance connections are over-represented (inset). Based on the fraction of axons with less-than expected variability, one can obtain the intersection point on the CV axis (dashed line, CV=0.54) and estimate the fraction of axon-dendrite pairs in the circuit that could have undergone Hebbian learning as at least 8.9%, whereas at least 35.1% show no sign of CV reduction possibly induced by Hebbian learning-related plasticity (inset). **(f-j)** Analysis of relation between highly consistent connections and average synapse size (f) in comparison to randomly drawn pairs of synapses from the entire synapse population (g) and in comparison to synapse pairs drawn randomly from the joint synapse population (h). Difference maps (i,j) show that in fact low-CV synapse pairs and larger synapse size are correlated (i); and that two populations of overrepresented low-CV synapses can be described (j, dashed areas); possibly corresponding to synapses with a history of LTP (up to 7.9%) and a small connection type or history of LTD (up to 15.5%).

We then measured the coefficient of variation (CV) of the axon-spine interface (ASI) area for random pairs of spine synapses (CV = 0.50 ± 0.32, mean ± s.d., n = 70,202) (Fig. 6e), for pairs of spine synapses sampled from the same dendrite but different input axons (CV = 0.50 ± 0.32; n = 64,934; indistinguishable from the random case, p = 0.07, one-sided Kolmogorov-Smirnov test), pairs of spine synapses sampled from the same axon but different target dendrites (CV = 0.49 ± 0.32; n = 54,256; slightly different from the random case, p = 1.7x10), and finally joint synapses established by the same axon onto the same dendrite (CV = 0.44 ± 0.30; n = 6,045; substantially smaller than the random case, p = 1.4x10^-40^). This data indicates that while heteronomous combinations of synapse pairs follow the random synapse pair distribution (Fig. 6e), for joint synapse pairs, significantly more pairs have substantially less synaptic variance (CV < 0.54) and substantially less pairs are found with a CV > 0.54. This data implies an excess of n=538 low-CV synapse pairs (i.e. 8.9%) compared to the random case (Fig. 6e), suggesting synapse pairs with a CV<0.54 as those that have been exposed to processes that homogenize synapse size. At least 35.1% (n=2,122) of connections involved in joint synapses show however a CV>0.54 that precludes such previous synapse size homogenization.

We next investigated the relationship between synapse size and its variability for joint synapse pairs, asking whether both the reduced size variability (Fig. 6e) and the average increase in joint synapse size (Fig. 6d) are caused by the same joint synapse pairs. For this we obtained the distribution of synapse size and its variability for all pairs of joint synapses (Fig. 6f) and subtracted a similar distribution randomly drawn from the size distribution of all (coupled and uncoupled) synapses (Fig. 6g). The resulting data clearly shows that reduced synapse size variability and increased synapse size are in fact correlated in the joint synapse pairs (Fig. 6i).

We finally wanted to investigate the relation between low-variability synapse pairs and their average size when controlling for the effect that joint synapses have an overall increased average synapse size (Fig. 6d). For this we again took the distribution of synapse size and its variability for all pairs of joint synapses (Fig. 6f), but this time we subtracted a similar distribution randomly drawn from the size distribution of only the joint synapses (Fig. 6h). The resulting data (Fig. 6j) again shows an enrichment of low-CV synapse pairs (Fig. 6j); Surprisingly, however, the data also indicates two separate enriched populations of synapse pairs with low CV: those with low CV and large synapse size (Fig. 6j, upper area, this range of synapse size variability and average synapse size contains 7.9% of all joint synapse pairs), and those with low CV and small synapse size (lower dashed area in Fig. 6j; 15.5% of all synapse pairs in this area).

Together, this data provides a connectomic fingerprint of the fraction of synapses in the circuit that have likely experienced previous homogenization of synapse size.

Both LTD (with saturation of synaptic weight decrease, (Egger et al., 1999)) and LTP are expected to yield such homogenization of synapse size. For at least about 24% of the joint synapse pairs (Fig. 6j), connectomic evidence for synapse size homogenization can be found. For at least about 35% of the joint synapses, however (Fig. 6e), synapse size homogenization cannot have occurred up to about one hour before the connectomic experiment (see (Bartol et al., 2015; Egger et al., 1999) for data on the time scales of synaptic weight change)).

## DISCUSSION

Using FocusEM, a set of tools for the semi-automated reconstruction of dense neuronal networks in the cerebral cortex, we have obtained the first dense circuit reconstruction from the mammalian cerebral cortex at a scale that allows the analysis of axonal rules of subcellular innervation-about 300 times larger than previous dense reconstructions from cortex (Kasthuri et al., 2015). We find that inhibitory axonal types specifically innervating certain postsynaptic subcellular compartments can be defined solely based on connectomic information (Figs 2,3); that in addition to inhibitory axons a fraction of excitatory axons exhibits such subcellular innervation preferences (Fig. 3); that the geometrical arrangement of axons and dendrites can explain only a moderate fraction of synaptic innervation, revoking random models of cortical wiring (Fig. 4); that a substantial thalamocortical synapse gradient in L4 gives rise to an enhanced heterogeneity of synaptic input composition at the level of single cortical dendrites (Fig. 5); and that the consistency of synapse size between pairs of axons and dendrites signifies fractions of the circuit with and without evidence for synaptic plasticity history, placing an upper bound on the “learned” fraction of the circuit (Fig. 6). Together, FocusEM allows the dense mapping of circuits in the cerebral cortex at a throughput that enables connectomic screening.

### Connectomic bounds on error rates

In the development of high-throughput reconstruction techniques for large-scale 3D-EM-based connectomics, the calibration of methodological progress using various definitions of error rates per cable length were initially important (Boergens et al., 2017; Cardona et al., 2012; Helmstaedter et al., 2011; Januszewski et al., 2018). With the ability to obtain dense connectomic maps ((Eichler et al., 2017; Helmstaedter et al., 2013; Wanner et al., 2016a) and this study), methodological proof-of-principle calibration can be replaced by a comparison of actual dense reconstructions and the required resources (Fig. 1p,q).

This becomes advantageous because error rates can be implicitly calibrated by the relevant connectomic analyses for a given dense reconstruction. In the presented data, we determined the required reconstruction accuracy by testing each finding for its sensitivity to the remaining reconstruction errors, and by performing manual control reconstructions where necessary (see Methods for details).

Once larger EM volumes are to be reconstructed, more investment in error rate reduction may be required (corresponding to higher manual annotation investments)-however, using the extensive labels from dense reconstructions as the present one, future reconstruction approaches may become already more efficient based on this previous knowledge alone.

### Connectomic evidence for synaptic plasticity

The extraction of connectomic evidence for synaptic size consistency requires further discussion. While the similarity of synaptic size has been previously used for arguments about synaptic plasticity and learning (Bartol et al., 2015; Bromer et al., 2018), an alternative source of decreased variance of synapse size for defined pairs of axons and dendrites is the establishment of consistently weaker or stronger synaptic connections between certain subtypes of pre-and postsynaptic neurons. While in the L4 circuit, the majority of excitatory connections is known to be established between local spiny neurons, and the observed synaptic size effects (Fig. 6) were also present when excluding TC inputs (data not shown), subtypes of excitatory neurons with different connectivity rules could contribute to the low-variance synaptic size regime. Our result on the lower bound of the fraction of the circuit that has no history of synaptic size homogenization (35%, Fig. 6e), is however unaffected by this cautionary notion.

### Outlook

The presented methods and results open the path to the consistent connectomic screening of mammalian tissue from various cortices, layers, species, developmental stages and disease conditions. The fact that even a small piece of mammalian cortical neuropil contains a high density of relevant information, so rich as to allow the extraction of possible connectomic signatures of the “learnedness” of the circuit, makes this approach a promising endeavor for the study of the structural setup of mammalian nervous systems.

### Author contributions

Conceived, initiated and supervised the project: MH; Performed experiments: KMB; Provided analysis methods: PH; developed and performed analyses: AM, MB, KMB, BS with contributions by all authors; wrote the manuscript: AM, MH with contributions by all authors.

## Acknowledgements

We thank Yunfeng Hua and Kristen Harris for discussions, Beth Cowgill for tissue staining and sample preparation, Elias Eulig, Robin Hesse and Matej Zecevic for contributions to automated neurite identification and soma reconstruction, Christian Guggenberger for management of the compute infrastructure, Dalila Rustemovic for support with figure preparation, and A. M. M. Al-Shaboti, M. S. E. A. Aly, C. Arras, N. Aydin, D. Baltissen, A. Bamberg, F. Y. Basoeki, N. Berghaus, A. Berghoff, L. Bezzenberger, N. M. Böffinger, S. M. Bohne, O. J. Brandt, A. B. Brandt, J. Buβ, L. Buxmann, D. E. Celik, H. Charif, N. O. Cipta, L. Decker, K. Desch, T. Engelmann, T. Ernst, J. Espino Martinez, T. A. R. Füller, D. J. Goffitzer, V. Gosch, J. C. Hartel, H. Hees, B. Heftrich, J. L. L. Heller, R. C. Hülse, O.-E. Ilea, R. Jakob, L. H. Janik, A. Jost Lopez, M. J. D. G. Jüngling, V. C. Kalbert, M. Karabel, A. Keβler, L. Kirchner, R. Kneiβl, T. Köcke, P. König, K. Krämer, F. K. Kramer, L. C. R. Kreppner, M. S. Kronawitter, J. Kubat, B. Kuhl, D. Kurt, C. Kurz, E. P. Laube, C. T. Lossnitzer, R. N. Lotz, L. A. Lutz, L. E. U. Matzner, S. Mehlmann, I. M. Meindl, I. Metz, M. Mittag, M.-L. Nonnenbroich, M. Nowak, N. Plath, A.-L. Possmayer, L. Präve, M. Präve, S. Reibeling, S. Reichel, K. J. P. Remmele, A. C. Rix, C. F. Sabatelli, D. J. Scheliu, N. Schmidt, L. M. Schütz, C. M. Schumm, A. K. Spohner, S. T. Stahl, B. L. Stiehl, K. T. Strahler, K. M. Trares, S. N. Umbach, S. S. Wehrheim, M. Werr, J. Winkelmeier, T. Winkelmeier, and S. Zimbelmann for webKnossos annotation work.

## METHODS

### Animal experiments

A wild-type (C57BL/6) male mouse was transcardially perfused at postnatal day 28 under isoflurane anesthesia using a solution of 2.5% paraformaldehyde and 1.25% glutaraldehyde (pH 7.4) following the protocol in (Briggman et al., 2011). All procedures followed the animal experiment regulations of the Max Planck Society and were approved by the local animal welfare authorities (Regierungspräsidien Oberbayern and Darmstadt).

### Tissue sampling and staining

The fixated brain was removed from the skull after 48h of fixation and sliced coronally using a vibratome. Two samples were extracted using a 1 mm biopsy punch (Integra Miltex, Plainsboro, NJ) from a 1 mm thick slice at 5 mm distance from the front of the brain targeted to layer 4 in somatosensory cortex of the right hemisphere. The corresponding tissue from the left hemisphere was further sliced into 70 μm-thick slices followed by cytochrome oxidase staining indicating the location of the coronal slice to be in barrel cortex.

Afterwards the extracted tissue was stained as in (Briggman et al., 2011). Briefly, the tissue was immersed in a reduced Osmium tetroxide solution (2% OsO_4_, 0.15 M CB, 2.5 M KFeCN) followed by a 1% Thiocarbohydrazide step and a 2% OsO_4_ step for amplification. After an overnight wash, the sample was further incubated with 1.5% Uranyl Acetate solution and a 0.02 M Lead(II) Nitrate solution. The sample was dehydrated with Propylenoxide and EtOH, embedded in Epon Hard (Serva Electrophoresis GmbH, Germany) and hardened for 48 h at 60 °C.

### 3D EM experiment

The embedded sample was placed on an aluminium stub and trimmed such that on all four sides of the sample the tissue was directly exposed. The sides of the sample were covered with gold in a sputter coater (Leica Microsystems, Wetzlar, Germany). Then, the sample was placed into a SBEM setup ((Denk and Horstmann, 2004), Magellan scanning electron microscope, FEI Company, Hillsboro, OR, equipped with a custom-built microtome courtesy of W Denk). The sample was oriented so that the radial cortex axis was in the cutting plane. The transition between L4 and L5A was identified in overview EM images by the sudden drop in soma density between the two layers (see Fig. 1b). A region of size 96 μm × 64 μm within L4 was selected for imaging using a 3 by 3 image mosaic, a pixel size of 11.24 × 11.24 nm^2^, image acquisition rate of 10 MHz, nominal beam current of 3.2 nA (thus a nominal electron dose of 15.8 e'/nm^2^), acceleration voltage of 2.5 kV and nominal cutting thickness of 28 nm. The effective data rate including overhead time spent during motor movements for cutting and tiling was 0.9 MB/s. 3,420 image planes were acquired, yielding 194 GB of data.

### Image alignment

After 3D EM dataset acquisition, all images were inspected manually and marked for imaging artifacts caused by debris present on the sample surface during imaging. Images with debris artifacts were replaced by the images at the same mosaic position from the previous or subsequent plane. First, rigid translation-only alignment was performed based on the procedures in (Briggman et al., 2011) which followed closely (Preibisch et al., 2009). The following modifications were applied: When shift vectors were obtained that yielded offsets of more than 100 pixels, these errors were iteratively corrected by manually reducing the weight of the corresponding entry in the least-square relaxation by a factor of 1000 until the highest remaining residual error was less than 10 pixels. Shift calculation of subsequent images in cutting direction was found to be the most reliable measurement and was therefore weighted 3-fold in the weighted least-square relaxation. The resulting shift vectors were applied (shift by integer voxel numbers) and the 3D image data was written in KNOSSOS format (Boergens et al., 2017; Helmstaedter et al., 2011).

### Previous usage of 3D image data

The 3D EM image dataset using the initial image alignment described above was previously utilized for methods development in (Berning et al., 2015), (Boergens et al., 2017) and (Staffler et al., 2017). Furthermore, reconstructions from this data were used as staining quality comparison in (Hua et al., 2015).

### Subimage alignment

To further improve the precision of image alignment, which we found to critically impact the quality of the automated volume segmentation, we performed the following steps. Each image of the raw dataset was cut into smaller images sized 256x256 pixels each. The offset calculation was run as described above (with the shift between neighboring subimages from the same original image set to zero). Additionally, we used a mask for blood vessels and nuclei (see below) to determine images which mostly contained blood vessels or somata. These images were assigned a decreased weight in the relaxation step. After the least-square relaxation, the shifts obtained for the subimages were used to create a smooth non-affine morphing of the original images, which were then exported to the 3D KNOSSOS format as above. All raw image data will be made available for inspection at demo.webknossos.org (see section on data availability).

### Methods description for software code

The following descriptions are aimed at pointing to the key algorithmic steps rather than enumerating all detailed computations.

### Blood vessel detection and correction for brightness gradient

Blood vessels were detected (Fig. 1e) by automated identification of regions of at least 0.125 μm^2^ with extreme brightness values (below 50 or above 162 at 8 bit depth) in each image plane, followed by manual inspection to exclude false positives. Image voxels within blood vessels were assigned the mean brightness of the entire dataset (mean=121). To correct brightness gradients across the image volume, the mean brightness was calculated for non-overlapping image blocks of 64 × 64 × 29 vx^3^, respectively, and the resulting marginal brightness distributions along the X, Y, and Z axes were smoothed and used to assign a multiplicative correction factor to each image block. The correction factor was linearly interpolated and multiplied to the brightness value of all non-blood vessel voxels within each of the image blocks.

### Nuclei and myelin detection

For the automated detection of nuclei and myelin, the following heuristics were applied. First, the voxel-wise brightness gradient was computed in the image data after smoothing by a 3-D kernel of size 21 × 21 × 9 vx and a standard deviation of ~33.5 nm. Nuclei were identified as regions of at least about 1.8 μm3 size with small brightness gradient and image brightness close to the mean image brightness. Myelin was detected as regions of low brightness sized at least about 0.35 μm3. Both nuclei and myelin detection were applied on overlapping image volumes of 912 × 912 × 416 vx^3^ size which were then truncated to non-overlapping volumes of 512 × 512 × 256 vx^3^ size.

### Volume segmentation using SegEM

To generate an initial automated volume segmentation, SegEM (Berning et al., 2015) was applied to image data cubes of size 1024 × 1024 × 512 vx^3^ with 256, 256, and 128 vx overlap along X, Y, and Z, respectively, using CNN 20130516T2040408,3 with parameters θ_ms_ = 10 vx and θ_hm_ = 0.25 (see Table 1 in (Berning et al., 2015)). At the edge of myelinated regions (see previous section), the CNN output was replaced with the minimum output value of -1.7 to enforce splits during the subsequent watershed-based volume segmentation.

### Segmentation neighborhood graph

For the determination of neurite continuity and synaptic interfaces (Fig. 1f), a segment neighborhood graph (region adjacency graph) was constructed in each of the non-overlapping segmentation cubes created by the SegEM step (see previous section). The neighborhood graph was constructed as in SynEM (Staffler et al., 2017). Briefly, two volume segments were called adjacent if there was a boundary voxel that contained both segments in its 26-neighborhood. The borders between adjacent segments were calculated as the connected components of all boundary voxels that had both segments in their 26-neighborhood. For each border, an edge between the corresponding segments was added to the neighborhood graph. The segment neighborhood graph is thus an undirected multigraph.

To extend the neighborhood graph beyond the non-overlapping segmentation cubes, pairs of segmentation cubes that shared a face in the x, y or z-direction were considered, and segments in the juxtaposed segmentation planes from the two segmentation cubes were matched if the number of matched voxels for a given pair of segments in the two planes was at least 10, and if the matched voxels constituted more than 90% of the area of the smaller segment. In these cases, an edge between the corresponding segments from the neighboring segmentation cubes was added to the neighborhood graph.

### Synapse detection with SynEM

For synapse detection, SynEM (Staffler et al., 2017) was applied to the segment neighborhood graph (see previous section) as in (Staffler et al., 2017). In brief, for each pair of adjacent volume segments, the subvolumes for SynEM feature aggregation (see (Staffler et al., 2017)) were determined by dilating the border between the two volume segments with spherical structuring elements of radius 40 nm, 80 nm and 160 nm, respectively, and then intersecting the dilated border the two adjacent volume segments, each. Interfaces with a border size of less than 151 voxels were discarded. Then, all interfaces in the segment neighborhood graph were classified using the SynEM classifier, yielding two SynEM scores for each interface, one for each of the two possible synapse directions.

In contrast to (Staffler et al., 2017), separate classifiers for interfaces onto spine segments (retrieved by TypeEM) and for all other interfaces were used. For interfaces onto spine segments, the classifier from (Staffler et al., 2017) was used. All interfaces onto spine segments with at least one score larger than -1.2292 according to the SynEM classifier (corresponding to 89% recall and 94% precision for spine synapses; see the test set of (Staffler et al., 2017)) were considered as synaptic interface candidates. For all other interfaces, a second classifier was trained using different training data and a different feature representation of interfaces. The training set of the second classifier consisted of all shaft and soma synapses of the SynEM training set and the shaft and soma synapses from two additional training volumes of size 5.75 × 5.75 × 7.17 μm^3^. The feature representation of interfaces for the second classifier consisted of all features of SynEM described in (Staffler et al., 2017), extended by four additional texture filter responses. The additional filter responses were voxel-wise probability maps for synaptic junctions, mitochondria, vesicle clouds and a background class obtained using a multi-class CNN. The CNN was trained on seven volumes of dense annotations for synaptic junctions, vesicle clouds and mitochondria (six volumes of size 3.37 × 3.37 × 3.36 μm^3^ that were also used for the methods comparison in (Staffler et al., 2017), and one additional volume of size 5.75 × 5.75 × 7.17 μm^3^) using the elektroNN framework (elektronn.org; see also (Dorkenwald et al., 2017)). Interfaces onto segments that were not classified as spines by TypeEM with at least one directional SynEM score larger than -1.28317 according to the second classifier (corresponding to 69% recall and 91% precision evaluated on all shaft synapses of the test set for inhibitory synapse detection of SynEM; see (Staffler et al., 2017)) were considered as synaptic interface candidates in addition to the synaptic interface candidates onto spines.

### ConnectEM classifier

To determine the continuity between adjacent volume segments (Fig. 1f), for each interface (see previous section) sized at least 10 vx, the SynEM filter bank and aggregation volumes (Staffler et al., 2017) were applied to the image and CNN output data, resulting in 6,426 texture-and 22 shape-features per interface. The features were used as input to an ensemble of 1,500 decision tree stumps trained with LogitBoost (Friedman et al., 2000) on 76,379 labeled edges obtained from proofread dense skeleton annotations of three (5 μm)^3^ cubes of neuropil. To adapt the SynEM interface classification method to a task on undirected edges, the ensemble was trained on both the forward and reverse direction of the labeled edges. Each edge in the segment neighborhood graph was then assigned a continuity probability by applying the classifier to the corresponding interface in random direction. Interfaces with less than 10 vx were treated as having a continuity probability of zero, edges across segmentation cubes were assigned a continuity probability of one.

### TypeEM classifier

To determine whether a volume segment belonged to a dendrite, an axon, or an astrocyte, and whether it was likely a dendritic spine head, we developed a set of four classifiers (”TypeEM”) as follows. Each volume segment was expanded into an agglomerate of up to 5 segments by iteratively adding the neighboring segment with the highest edge continuity probability to the agglomerate. Agglomeration was restricted to the subgraph induced by the edges with at least 92% continuity probability to prevent merge errors. Then, the following set of features was computed for the agglomerates: 918 texture features from the SynEM filter bank (Staffler et al., 2017) applied to the image and CNN output data and pooled over the segment agglomerate volume; 6 shape features as in SynEM; the 0th-to 2nd-order statistical moments of the agglomerate volume; the eigendecomposition of the 2^nd^ order statistical moment; the 0^th^-to-2^nd^ order statistical moments of the surface of the agglomerate after rotation of the agglomerate to the principal component of all its voxels; same as before but for the convex hull of the agglomerate; volume-to-surface area ratio, compactness (i.e., (surface area)^3^ / volume^2^), clusters of normal unit vectors, hull crumpliness and packing (Corney et al., 2002); estimates of the distance (Osada et al., 2001) and thickness (Yi et al., 2004) histograms from sampling random point pairs on the agglomerate’s surface.

This yielded a total of 1,207 shape features and 924 SynEM features; these were then taken as input to an ensemble of 1,500 decision tree stubs trained using LogitBoost (Friedman et al., 2000). 14,657 training samples were obtained by one expert (AM) marking all spine head segments and assigning the neurite / glia type to each process in three densely reconstructed (5 μm)^3^ cubes of neuropil (same as in previous section “ConnectEM classifier”).

Together, these data were used to train one-versus-all TypeEM classifiers for axons, dendrites, and astrocytes. The classifiers reached the following classification performance on a separate test cube sized (5 μm)^3^: Axon classifier: 91.8% precision (P) and 92.9% recall (R); dendrite classifier: 95.3% P, 90.7% R; astrocyte classifier: 97.2% P, 85.9% R (at maximum area under precision-recall curve). The spine head classifier was trained on a feature set calculated as above, with the exception that the agglomeration step was omitted, and achieved 92.6% P and 94.4% R.

For subsequent processing, the TypeEM classifier scores were transformed to probabilities using Platt scaling (Platt, 1999. Probabilistic Outputs for Support Vector Machines and Comparisons to Regularized Likelihood Methods. In: Advances in large margin classifiers.). Finally, the one-versus-all axon, dendrite, and astrocyte probabilities of each segment were combined to multi-class probabilities by rescaling them by the inverse of their sum.

### Automated reconstruction of dendrites

For the reconstruction of dendrites, we first selected all SegEM segments with a TypeEM dendrite probability (Fig. 1h) of at least 0.3 and a volume of at least 500 vx. In the subgraph induced by these segments we deleted all edges that corresponded to an interface of less than 300 vx size or a neurite continuity probability below 98%. The graph was then used to cluster the dendritic segments into connected components, yielding dendrite agglomerates.

To reduce the effect of TypeEM misclassifications, we used the fraction of myelinated surface area to remove agglomerates from the dendrite class (calibrated based on 50 random agglomerates with a myelinated surface fraction between 0.05 and 0.5): Agglomerates had a total volume of at least 200,000 voxels, had a myelinated surface fraction above 0.25 (or above 0.08 if the agglomerate comprised more than 25 segments); agglomerates did not contain somatic segments.

The myelinated surface fraction was calculated for each agglomerate with a total volume of above 5 μm^3^. All neighboring segments of the agglomerate were identified according to the neighborhood graph, and the area of interfaces onto neighboring myelin segments, defined as having at least 50% of their volume intersecting with the myelin heuristic, were added up. This area was then divided by the total area of all interfaces between the agglomerate and other segments.

### Reconstruction of cell bodies (somata)

Cell bodies were reconstructed from the volume segmentation of each cell’s nucleus (see above). First, we identified all SegEM segments which were contained in a nucleus with at least 50% of their volume. Then we added all direct neighbors of these segments according to the neighborhood graph. Then we iteratively extended the soma volumes along the neighborhood graph with the following constraints: only consider segments with a size of at least 2,000 voxels and a center of mass at a maximal distance of 8 μm from the center of mass of the corresponding nucleus; only consider edges in the neighborhood graph with a continuity score above 0.98; do not consider edges if the segments’ vessel score or its myelin score were above 0.5. Then, all connected components of segments that were completely enclosed by soma segments according to the neighborhood graph were added to the respective soma. Finally, all segments with more than 80% of their surface adjacent to soma segments were added iteratively (10 iterations).

### Soma-seeded reconstruction of neurons

For the reconstruction of those neurons that had their cell body in the dataset (n=89 with dendrites reconstructed in the dataset, Fig. 1l,m, Suppl. Material 1), all dendrite agglomerates from the automated dendrite reconstruction that overlapped with a given soma volume (see previous section) were combined into one agglomerate for each of the neurons.

### Iterative semi-automated correction of whole cell agglomerates

The remaining errors in the soma-seeded neuron reconstructions (see previous section) were corrected in a semi-automated procedure that consumed 9.7 hours for all neurons, i.e. 5.18 minutes per neuron. Soma-based neuron reconstructions were inspected for merger errors in the 3D view of webKnossos, and mergers were corrected by deletion of nodes in the neighborhood graph of the neuron reconstruction. Then, endings of the neuron were detected (see below), and reconstructions at the endings were performed in webKnossos until a dendrite agglomerate was reached that was obtained from the automated dendrite reconstruction (see previous section). The inspection for mergers and the detection of endings in the dataset was iterated until only real endings or endings at the dataset boundary were left.

### Automated axon reconstruction

For the reconstruction of axons, we first selected all SegEM segments with a volume of at least 300 vx and a TypeEM axon probability (Fig. 1h) of at least 50%. The subgraph induced by these segments was partitioned into connected components (axon agglomerates) after removal of edges corresponding to interfaces with less than 60 vx size or with a neurite continuity probability below 97%. Next, for each segment that was part of an axon agglomerate, we computed the first principal component of its voxel locations and used its degree of variance explanation as an indicator for the directedness of the segment. We then determined for each interface between the agglomerate’s segments and all neighboring segments the alignment of the interface’s normal vector with the segment direction. Based on this, we obtained an ending score for each interface of the segment, and at locations with high scores, the axon agglomerate was grown into neighboring segments under the following additional constraints: the neighboring segment had an axon probability of at least 30%; the interface had a size of at least 40 vx; the neighborhood graph edge had a neurite continuity probability of at least 80%. This growth process was repeated ten times.

Finally, we compensated for the heightened rate of merge errors in proximity to the dataset boundary that results from decreased alignment quality. Edges that were closer than 2 μm to dataset boundary and had a neurite continuity probability below 98% were removed from the axon agglomerates.

Then, all axon agglomerates of length 5 μm and above were used for the following focused annotation steps. Length of agglomerates was computed as the summed Euclidean length of all edges in the minimal spanning tree of the center of masses of the agglomerate’s segments.

### FocusEM ending detection and query generation

To determine the endings of axons at which focused annotation could be seeded, we used the following procedure. For each segment in an axon agglomerate we took the segments that are direct graph neighbors or that come within 1 μm distance, and computed the first principal component of their volume. We then identified all segments where the principal component of the local surround explains at least 50% of the variance, and determined the borders on the axon agglomerate surface that were aligned to that axis (i.e., all interfaces for which the vectors from the center of mass of the local surround to the center of mass of the interface were at an angle of at most cos^-1^(0.8) ≈ 37 °). Finally, the identified interfaces were grouped using a cutoff distance of 600 nm and reduced to the interface best aligned to the surround’s principal component. We determined the point within 500 nm of each interface that is closest to the core of the axon agglomerate and used it together with the principal component of the local surround as the start position and orientation of a focused annotation query in webKnossos. Interfaces within 3 μm of the dataset boundary were excluded from query generation.

FocusEM axon queries were performed in webKnossos flight mode. The volume map of all axon agglomerates larger than 5 μm was used to dynamically terminate flight paths when a user entered already reconstructed agglomerates (this was implemented in a custom script using the webKnossos frontend APITo reduce the delay between subsequent queries, we implemented a “hot switching mode” in webKnossos such that the next query was already loaded in the background while answering the current query. With this, an immediate switching (amounting to a jumping to query locations in the dataset) was possible that yielded negligible lag between tasks.

### Query analysis

FocusEM queries yielded linear skeletons from webKnossos flight mode. For each node of a given skeleton we determined the overlap with axon agglomerates in the (3 vx)^3^ cube around each skeleton node (a skeleton was considered to overlap with an axon agglomerate if the agglomerate was contained in at least 54 vx around the skeleton nodes). For the overlapping agglomerates, we determined the corresponding agglomerate endings within 300 nm distance from the skeleton nodes. Based on the configuration of agglomerate overlaps, agglomerate endings reached by the queries and proximity of the query to the dataset boundary, the query results were either accepted as is, re-queried or discarded (see code files below for detailed decision tree). When locations were queried multiple times, the information on agglomerate and ending overlap was used to keep only minimal subsets of skeleton tracings for the final axon agglomerates (see “Iteration between ending and chiasma detection”). For connectome analysis and display, volume segments that had not yet been assigned to any axon agglomerate and that overlapped with the user skeleton from the flight mode queries were collected and added to the agglomerate volume.

### Chiasma detection and queries

To identify mergers, we detected geometric configurations (Fig. 1k) with more than two-fold neurite crossings after agglomeration. Chiasmata were detected by counting the number of intersections of the graph representation of a given agglomerate with a sphere centered on the nodes of the graph. For this, the agglomerate was reduced to the connected component contained within a sphere of 10 μm radius around the current node, and then all edges within a sphere of radius of 1 μm were removed. The remaining graph components were considered sphere exits. If four or more sphere exits were found, the node at the sphere center was labeled as a chiasmatic node. Within axon agglomerates, the chiasmatic nodes were clustered using a cutoff distance of 2 μm and subsequently reduced to the node closest to the center of mass of the cluster. At these locations, queries from the sphere exits pointing towards the sphere center were generated and annotated as described for the ending queries (Fig. 1n,o). The webKnossos flight mode annotations of chiasma queries were stopped when the annotator left the bounding box around all exit locations.

### Chiasma query interpretation

To decide which of the exits contributing to a given chiasma should remain connected and which should be disconnected, we used the query results from all chiasma exits. The full set of results enabled the detection of chiasmata with contradictory query answers, partial automated error correction, and the re-querying of a minimal set of exits. Chiasmata with a full and contradiction-free set of answers were solved by removing the edges within the center 1μm sphere from the agglomerate mst and by subsequent reconnection of the exits based on a minimal set of flight queries.

### Iteration between ending and chiasma detection

Following automated axon reconstruction, the FocusEM queries for ending and chiasma annotations were applied iteratively.

### Spine head agglomeration

Spine heads were agglomerated by connecting neighboring segments with a TypeEM spine head probability above 50% that were connected by an edge with neurite continuity probability of at least 98%. Spine head detections within blood vessels were discarded. This yielded 415,797 spine head agglomerates.

### Spine attachment

Of the 415,797 spine head agglomerates, 5.6% got attached to a dendritic shaft during automated dendrite agglomeration (see above). We then implemented a greedy walk strategy from spine heads to the corresponding dendritic shafts. The walk was terminated upon reaching a dendrite agglomerate of at least ~1.1 μm^3^(10^5.5^ vx) and was restricted to at most ten steps along continuity edges, each having a neurite continuity probability of 25% or more and only involving segments with axon probability below 80%. With this, an additional 206,546 (49.7%) spine heads could be attached to the corresponding dendrite. For the remaining spines, SegEM mergers in the very thin spin necks typically prevented the spine attachment heuristics to be successful. We instead seeded manual annotation in the 164’969 remaining spine heads with a distance of at least 3 μm from the dataset boundary, asking annotators to connect these to the dendritic shafts. This consumed 900 work hours total and resulted in a final spine head recall of 88.6% for spine heads further than 10 μm from the dataset boundary.

### Synapse agglomeration

Synaptic interface candidates (n = 864,405 out of which 605,569 were onto spine segments and 258,836 onto other segments) detected by interface classification were discarded if the score both synapse scores were larger than -2 or the continuity probability of the corresponding edge was larger than 0.7, or the myelin score of the pre-or postsynaptic segment was larger than 0.375 or the presynaptic segment was contained in the soma volumes. The remaining synaptic interface candidates (n= 862,350) were restricted to those with a center of mass more than 3 μm from the segmentation volume boundary (yielding n=696,149 synaptic interfaces with a pre- and postsynaptic segmentation object, each, that are used in the following analyses).

To consolidate synaptic interfaces, the following steps were applied: First, all presynaptic segmentation objects contributing to any of the synaptic interfaces were combined, if they were connected to each other by at most two steps on the segmentation neighborhood graph with each step along an edge above 0.98 ConnectEM score. The same was applied to all postsynaptic segmentation objects. Then, all synaptic interfaces between the combined pre- and postsynaptic segmentation objects were combined into one synapse, each. Synapse agglomerates for which at least one postsynaptic segment was part of the spine head agglomerates were considered as spine synapse agglomerates. A spine synapse agglomerate was called “primary spine innervation” if it contained the interface with the highest SynEM score onto a given spine head agglomerate, and “secondary spine innervation” otherwise. Multiple synapse agglomerates between an axon agglomerate and a spine head agglomerate were merged into a single synapse agglomerate. The center of mass for a synapse agglomerate was calculated as the component-wise mean of the centers of mass of the individual interfaces. The area of a synapse agglomerate was calculated as sum of the border areas of the individual interfaces.

### Soma synapse exclusion heuristic

Synapse agglomerates for which at least one postsynaptic segment was part of the soma agglomerates were considered as soma synapses. Synapse agglomerates were clustered based on their center of mass using hierarchical clustering with single linkage and a distance cutoff of 1 μm. If a synapse agglomerate cluster contained a spine synapse which was the only synapse onto the corresponding spine head agglomerate, then all soma synapses of the synapse agglomerate cluster were discarded. Synapses from excitatory axons onto somata of spiny cells were ignored for the analysis of subcellular target specificity and geometric predictability.

### Connectome aggregation

The connectome was constructed using the axon agglomerates, postsynaptic agglomerates (dendrites, somata and axon initial segments), and synapse agglomerates. For each pair of an axon and postsynaptic agglomerate, all synapse agglomerates that had a presynaptic segment in the axon agglomerate and a postsynaptic segment in the postsynaptic agglomerate were extracted and associated with the corresponding axon-target connection. The total number of synapses of a connection was defined as the number of synapse agglomerates associated with that connection. The total border area of a connection was defined as the sum of the border area of all synapse agglomerates. All of the following analyses were restricted to axons with at least ten output synapses.

### Target Class Detection Heuristics

To determine the post-synaptic target classes apical dendrites (AD), smooth dendrites (interneuron dendrites, SD), axon initial segments (AIS), proximal dendrites (PD) and cell bodies (SOM), the following heuristics were used: Cell bodies were identified based on the detection of nuclei as described in “Reconstruction of cell bodies”. The non-somatic postsynaptic components of the soma-based neuron reconstructions were marked as proximal dendrites. Smooth dendrites were identified by having a spine rate (i.e., number of spines per dendritic trunk path length) below 0.4 per μm (Kawaguchi, Karuba, Kubota, 2006), Fig. 2d, unless identified as apical dendrites. For the analysis of target class specificities and geometric predictability, the dendrites of soma-based interneuron reconstructions were considered as smooth, but not proximal dendrites.

For the identification of apical dendrites, all dendrite agglomerates that intersected with the pia-and white matter-oriented faces of the dataset were manually inspected in webKnossos (total of 422 candidates, total inspection time 5 hours for an expert annotator) with the inspection criteria: directed trajectory along the cortical axis; maximally two oblique dendrites leaving the main dendrite; spine rate of non-stubby spines of at least about one every two micrometers.

Contradictory class assignments between SD and AD occurred for 46 dendrites and were resolved by manual inspection in webKnossos.

The axonal part of soma-based neuron reconstructions which was more proximal than the first branch point was considered as axon initial segment. Vertically oriented agglomerates that entered the dataset from the pia-end of the dataset and had no spines or output synapses, and transitioned into a clearly axonal process closer to the white matter boundary of the dataset were also identified as axon initial segments.

### Definition of inhibitory and excitatory axons

Inhibitory and excitatory axons were separated based on the fraction of their synapses marked as primary spine innervations (see “synapse agglomeration”) (Fig. 3a). To automatically resolve remaining merge errors between these two axon classes, we split axons between the two modes of the spine rate distribution (20 to 50% of synapses being primary spine innervations) at all their branch points (see “Chiasma detection and queries”). Then, we defined excitatory axons as those with more than 50% of synapses being primary spine innervations and termed axons with less than 20% primary spine innervations inhibitory.

### Definition of thalamocortical axons

To identify those excitatory axons that were likely originating from the thalamus we used the fact that thalamic axons from VPM have been described to establish large multi-synaptic boutons at high frequency in mouse S1 cortex ((Bopp et al., 2017); see Fig. 3h,i). We quantitatively applied these criteria by measuring the density of primary spine innervations (PSI) per axonal path length, the average number of PSI per axonal bouton, the fraction of axonal boutons with multiple PSIs, and the median bouton volume. Boutons were defined as clusters of PSI with an axonal path length of less than 2.4 μm between the cluster centers. In a calibration set of ten manually identified corticocortical and ten thalamocortical axons these features were discriminatory. We combined them into a single thalamocortical axon probability using logistic regression. Excitatory axons with a TC probability of at least 60% were identified as thalamocortical.

### Subcellular specificity analysis

First, we assumed that all synapses of a given axon class have the same probability to innervate a particular postsynaptic target class (as above). We then inferred this first-order innervation rate for each axon-and postsynaptic target-class by searching for the probability which best explains whether or not an axon innervated the target class under a binomial model. The optimized binomial model was then used together with the measured number of synapses of each axon to calculate the expected distribution of target innervation rates. A one-sided Kolmogorov-Smirnov was used to test for the existence of a subpopulation with increased target innervation rate. To identify those axons that innervated a given target class beyond chance (Fig. 3k), we computed the probability *p^(t)^ _meas,i,k_* of finding at least the measured fraction of synapses onto target *t* for each axon *i* from axon class k. The p-values were also calculated for the expected distribution of target innervation rates and combined with *p^(t)^meas,i,k* to estimate the p-value threshold p∩^t^_k_ at which the false discovery rate q (Storey and Tibshirani, 2003) crosses 20%. 80% of the axons with *p^(t)^_meas,i,k_* < *p∩^(t)^_k_* are innervating target *t* with a rate above the first-order innervation probability and are thus called to be t-specific.

For the analysis of 2^nd^ order innervation specificity (Fig. 3l,m), we reported the fraction of synapses onto target *τ* by t-specific axons of class *k* after removal of synapses onto t. This innervation rate was compared against the fraction of synapses onto target *τ* by all axons of class k.

### Geometrical predictability analysis

To determine whether the found innervations can be predicted by geometrical measurements we used the following model: For each axon we determined the total surface area of the target classes that were contained within the cylinder of radius *r_pred_* around the axon (Fig. 4a-c) and compared it to the actually innervated target fraction of each axon (Fig. 4c,d). We then analyzed the correlation between the availability of the target surfaces and the actually established synapses on these target classes (Fig. 4e).

To obtain an overall predictability quantification, we then computed the coefficient of determination (R^2^) using the following model: For all axons of given type, we used the fraction of target innervations and fractional surface availabilities in a given surround of radius r_pred_ to find the optimal multivariate linear regression parameters. To estimate best-case geometric predictability, we then calculated the R^2^ value as 1 minus the ratio of the residuals to synaptic variance on the same axons used for parameter optimization, while correcting for the variance introduced by the finite number of synapses per axon. Accordingly, we used the axons’ fractional surface availabilities within r_pred_ and absolute synapse numbers to calculate the expected binomial variance, and subtracted it from the squared residuals. If the remaining squared residual of an axon was negative after correction, it was set to zero.

This analysis made several assumptions that were in favor of a geometrical explanation of synaptic innervation (therefore the conclusions about a minimal predictability (Fig. 4f) are still upper bound estimates): it was assumed that the number of synapses for a given axon was already known; in most settings, only average synapse rates are known for a given circuit; it also assumed that a precise knowledge of the axonal trajectory and the surrounding target surface fractions were available; again, this is usually only available as an average on the scale of rpred of several 10’s of micrometers.

To relax the assumption of complete knowledge about target availabilities, we repeated the above R^2^ analysis for a model in which the predicted fractional innervation of a target is the fractional surface availability of that target.

### Synaptic input/output maps and spatial synapse distributions

To determine the spatial distribution of synaptic inputs along dendrites, we used the soma-based neuron reconstructions and determined for each synapses onto their dendrites the shortest pairwise path length between all somatic and postsynaptic segments (Fig. 5a,b). Input synapses were assigned the class of the corresponding axon (as described above) and then pooled over all neurons. Synapses originating from axons of unknown type (e.g., axon had less than ten synapses and thus was excluded from classification) were ignored. The spatial output map was derived from axon tracings, which were seeded in all somata and then mapped onto the segmentation (see “Query analysis”) to find output synapses based on segment overlap (Fig. 5c,d).

The spatial synapse distribution of a given axon class was obtained by projecting the center of mass of the corresponding synapses (see “Synapse agglomeration”) onto the XY plane, where × is the pia-white matter axis, and then calculating a kernel density estimate thereof (Fig. 5e). The synapse ratio along the cortical axis was computed as the ratio of absolute synapse counts per histogram bin (Fig. 5f-h).

To quantify the effect of soma location on synaptic innervation, we calculated for each soma-based neuron reconstruction the center of mass of all somatic segments, and the fraction of excitatory input synapses that originate from thalamocortical axons. We performed multivariate linear regression in the YZ plane orthogonal to the cortical axis and corrected the measured synapse fraction before quantifying the effect of cortical depth using univariate linear regression.

Finally, the soma-based neuron reconstructions were manually split into their primary dendrites to assess the effect of dendritic orientation on synaptic inputs. The orientation of a dendrite was calculated as the volume-weighted mean of the unit vectors from the soma (as above) to the center of mass of the corresponding SegEM segments. Finally, the dot product *dp* of the resulting vector (after renormalization) with the unit vector along the cortical axis was put in relation to the ratio of the dendritic synapse fraction to the synapse fraction of the corresponding neuron. The linear regression of these two quantities was evaluated based on the coefficient of determination, whereas the pia-*(dp* <-0.5) and white matter-oriented dendrites *(dp* > 0.5) were compared based on a two-sample t-test.

### Synapse-size consistency analysis

To determine the consistency of primary spine synapses between a given axon-dendrite pair, we calculated the axon-spine interface area (ASI, (de Vivo et al., 2017) (Staffler et al., 2017)) of a synapse as the total contact area between the corresponding axon and spine head agglomerates. For axon-dendrite pairs connected by exactly two primary spine synapses, we then calculated the coefficient of variation (CV) of the ASI areas by *CV = 2^1/2^ (ASII-AS2) /(ASH + AS2)* with *ASII* and ASI_2_ being the larger and smaller of the two ASI areas, respectively. To avoid false same-axon same-dendrite (AADD) pairs caused by remaining merge errors in the axon reconstruction, this analysis was performed only after splitting all axons at all branch points. The measured distribution of CV values was compared against the CV values obtained by randomly drawing pairs from the observed ASI area distribution (Fig. 6e,f). To test whether same-axon same-dendrite (AADD) primary spine synapse pairs have a lower CV than random pairs, a one-sided Kolmogorov-Smirnov test was used. The critical CV value that defines the upper limit of the range in which AADD pairs occur more often than expected was determined by searching for the intersection of the kernel density estimates of the observed and expected CV distributions. A lower bound on the fraction of overly-consistent primary spine synapse pairs is given by the difference between the cumulative probability functions of the two distributions at the critical CV value. Finally, we built kernel density estimates of the probability density function over the two-dimensional space defined by the CV and mean log-io(ASI) for AADD pairs, random pairs, and random pairs of AADD synapses. The differences between these density estimates were used to extract the non-random components in CV-ASI relationship of AADD pairs relative to random pairs and random pairs of AADD synapses, respectively.

### Comparison between dense reconstructions

For the comparison of published dense reconstructions and the invested resources (Fig. 1p,q), we used the following numbers: Dense reconstruction in the mouse retina (Helmstaedter et al., 2013): about 640 mm reconstructed neuronal path length, about 20,000 invested work hours; Dense reconstruction in the mouse cerebral cortex (Kasthuri et al., 2015): 6.75 mm of path length reconstructed within 253 hours (37.5 h/mm, 4.5 μm/μm^3^, 1500 μm^3^ reconstructed; see (Berning et al., 2015) for derivation of numbers); Dense reconstruction in the zebrafish olfactory bulb (Wanner et al., 2016b): 492 mm path length with 25,478 invested work hours; Dense reconstruction in the fly larval nervous system (Eichler et al., 2017): 2.07 meters (based on skeleton reconstructions in Supplement of (Eichler et al., 2017)) with 28,400 hours investment (73 μm / h; see (Schneider-Mizell et al., 2016)); L4 dense reconstruction: 2.724 meters (of these in mm: dendritic shafts 342, dendritic spines 551, dendrites connected to a cell body in the volume 62.5, axons connected to a cell body in the volume 6.5; axons 1760; note about 80% of the volume is dense neuropil) within 3,982 hours.

### Computational cost estimate

For the estimation of the total computational cost, a runtime of 5 hours for SynEM, 72 hours for TypeEM and 24 hours for all other routines on a cluster with 24 nodes each with 16 CPU cores and 16 GB RAM per core was used. The runtime was converted to resources using 0.105 USD/h per CPU core with 16 GB RAM (Amazon EC2: 6.7 USD/h for 64 CPU cores with 1000 GB RAM). The computational cost for Flood-Filling Networks was calculated using 1000 GPUs that ran for a total wall time of 16.02 hours (Suppl. Table 3, (Januszewski et al., 2018)) and a cost of 0.9 USD/h for a single GPU which was multiplied by the ratio of the sizes of our dataset (61 × 94 × 92 μm^3^) and the dataset used in (Januszewski et al., 2018) (96 × 98 × 114 μm^3^).

### Error Measurements

To quantify the errors remaining after axon reconstruction, we chose the same 10 randomly selected axons (total path length, 1.72 mm) that had also been used for error rate quantification in (Boergens et al., 2017). These axons were not part of any training or validation set in the development of FocusEM. Repeating the analysis described in (Boergens et al., 2017) for the largest axon agglomerate overlapping with the ground truth axon, respectively, yielded a total number of 22 errors, of which 15 were continuity errors (compare to panels 1l,m in (Boergens et al., 2017)).

The error rates and recall of soma-based dendrite reconstructions were calculated from proofread ground truth annotations comprising a total of 89 cells and 64.08 mm path length. Each node of the ground truth skeleton was marked as recalled if it overlapped with the corresponding dendrite agglomerate (see “Query analysis”), or flagged invalid if placed outside the segmented volume. A ground truth edge was considered recalled if both end nodes were recalled, or invalid if any of the end nodes was invalid. 54.51 mm, or 87.3%, of the 62.46 mm valid ground truth path length were recalled. Split errors, by definition, result in partial dendrite reconstructions and were thus detected as non-recalled ground truth fragments with at least 5 μm path length. The detected were proofread, yielding a total of 37 split errors.

The identification of axonal target specificity (Fig. 3) was insensitive to split-and merger errors, because as long as axonal reconstructions were long enough to provide meaningful statistical power for the analyses, split axons were expected to correctly sample target specificities and mergers of axons were expected to only dilute specificities. Therefore the results about the existence of target specific inhibitory and excitatory axon classes represent a lower bound of specific wiring. The results on the lack of geometric predictability (Fig. 4) were similarly unaffected by remaining split and merge errors.

For the finding that no inhibitory axons show target specificity for AIS in L4 (Fig. 3i), however, we needed to control that this lack of specificity was not induced by remaining axonal merge errors. We manually inspected a subset of 10 axons innervating AIS. Only one synapse (out of more than 100 synapses) was erroneously added to an AIS innervating axon due to a merger, thus providing no evidence that the lack of AIS target specificity could be an artifact of merged axons.

For the results on synaptic input composition (Fig. 5) we varied the sensitivity of our detection of TC axons and found that also for detections with a higher TC axon recall and a lower recall at higher precision the conclusions were unchanged.

The results on synaptic size consistency could be strongly affected by the remaining merge errors in axons, diluting data on consistent synapses when merging unrelated axons together. To control for this, we obtained the results in Fig. 6 using axons for which all 3-fold intersections in all axons had been artificially split before the analysis. For the results in Fig. 3, we repeated analyses after splitting of axons and found the key conclusions unaltered.

### Statistical methods

The following statistical tests were performed (in order of presentation in the figures):

The existence of axon subpopulation with unexpectedly high synapse rate onto a given target class was tested using the one-sided Kolmogorov-Smirnov test (Fig. 3i,j). Axons belonging to a given target-specificity class were identified based on the false detection rate criterion (q=20%, (Storey and Tibshirani, 2003)) (Fig. 3k).

The degree to which synaptic variance is explainable by geometry-based models was evaluated using the coefficient of determination (R^2^) (Fig. 4f). Binomial variance was corrected for by subtracting the surface fraction-based expected binomial variance from the squared residuals. The result was set to zero, if negative.

Path-length dependent axonal synapse sorting was tested using a two-sided t-test. F-tests were used to evaluate synaptic gradients as function of cortical depth (Fig. 5f,h) or dendritic orientation (Fig. 5j).

To test whether the axon-spine interface areas of a given spine synapse pair configuration were more similar than randomly sampled pairs, a one-sided Kolmogorov-Smirnov test was used (Fig. 6e).

### Data availability, software availability

All raw segmentation data, skeleton annotations, connectomic data, and software developed and used in this study will be made publicly available upon publication.

